# Neuregulin1 nuclear signaling influences adult neurogenesis and regulates a schizophrenia susceptibility gene network within the mouse dentate gyrus

**DOI:** 10.1101/2022.08.10.503469

**Authors:** Prithviraj Rajebhosale, Alice Jone, Kory R. Johnson, Rohan Hofland, Camille Palarpalar, Samara Khan, Lorna W. Role, David A. Talmage

**Author notes:** Corresponding Author: David Talmage. Co-first authors (Prithviraj Rajebhosale, Alice Jone). These authors contributed equally (Camille Palarpalar, Samara Khan).

## Abstract

Neuregulin1 (Nrg1) signaling is critical for aspects of neuronal development and function from fate specification to synaptic plasticity. Type III Nrg1 is a synaptic protein which engages in bi-directional signaling with its receptor ErbB4. Forward signaling engages ErbB4 phosphorylation, whereas back signaling engages two known mechanisms: 1. local axonal PI3K-AKT signaling, and 2. cleavage by gamma secretase resulting in cytosolic release of the intracellular domain (ICD), which can traffic to the nucleus (Bao, Wolpowitz et al. 2003, Hancock, Canetta et al. 2008). To dissect the contribution of these alternate signaling strategies to neuronal development we generated a transgenic mouse with a missense mutation (V_321_L) in the Nrg1 transmembrane domain that disrupts nuclear back signaling with minimal effects on forward signaling or local back-signaling and was previously found to be associated with psychosis (Walss-Bass, Liu et al. 2006). We combined RNA sequencing, retroviral fate mapping of neural stem cells, behavioral analyses, and various network analyses of transcriptomic data to investigate the effect of disrupting Nrg1 nuclear back-signaling in the dentate gyrus (DG) of male and female mice.

The V_321_L mutation impairs nuclear translocation of the Nrg1 ICD and alters gene expression in the DG. V_321_L mice show reduced stem cell proliferation, altered cell cycle dynamics, fate specification defects, and dendritic dysmorphogenesis. Orthologs of known schizophrenia (SCZ)-susceptibility genes were dysregulated in the V_321_L DG. These genes coordinated a larger network with other dysregulated genes. WGCNA and protein-interaction network analyses revealed striking similarity between DG transcriptomes of V_321_L mouse and humans with schizophrenia.

**SIGNIFICANCE STATEMENT:** Synaptic contact is predicted to be a regulator of the generation of nuclear signaling by Nrg1. Here we show that a schizophrenia-associated mutation in Nrg1 disrupts its ability to communicate extracellular signals to the neuronal genome which results in altered expression of a gene network enriched for orthologs of schizophrenia-susceptibility genes. The striking overlap in functional and molecular alterations between a single rare homozygous missense mutation (V_321_L) and schizophrenia patient data (complex polygenic and environmental burden) underscores potential convergence of rare and common variants on the same cellular and molecular phenotypes. Furthermore, our data indicate that the evolutionarily conserved gene networks that form the basis for this risk are necessary for coordinating neurodevelopmental events in the DG.

## INTRODUCTION

The Neuregulin 1 (Nrg1)-ErbB4 signaling axis is an important regulator of synapse formation and function throughout the nervous system (Harrison and Weinberger 2005, Walsh, McClellan et al. 2008, Agarwal, Zhang et al. 2014). Type III Nrg1, located pre-synaptically, participates in bidirectional signaling i.e., acting as both a ligand and a receptor for ErbB4 (Bao, Wolpowitz et al. 2003, Talmage 2008). Upon binding ErbB4, the Type III Nrg1 C-terminal intracellular domain (ICD) can be liberated into the cytosol from the membrane by an intramembrane proteolytic event mediated by the gamma secretase enzyme complex. Following this cleavage, the ICD translocates to the nucleus where it can function as a transcriptional regulator (nuclear back signaling) (Bao, Wolpowitz et al. 2003). Additionally, binding of ErbB4 can also induce local activation of PI3K-AKT signaling (Hancock, Canetta et al. 2008). Thus, activation of Type III Nrg1 by ErbB4 can induce two distinct modes of signaling. Cortical neurons cultured from Type III Nrg1 KO mice show deficits in axonal and dendritic outgrowth and branching, which were rescued by re-expression of full-length Type III Nrg1 protein (Chen, Hancock et al. 2010). However, nuclear back signaling defective forms of Type III Nrg1, lacking the nuclear localization sequence (NLS) or mutations in the gamma secretase cleavage site, were only able to rescue axonal growth and not dendritic growth (Chen, Hancock et al. 2010). In line with this, Type III Nrg1 hypomorphic mice showed reduced hippocampal dendritic spine density. Overexpression of the nuclear Nrg1 ICD resulted in increased dendritic spine density in cultured hippocampal neurons while overexpression of Nrg1 ICD lacking the NLS did not (Chen, Hancock et al. 2010, Fazzari, Snellinx et al. 2014). Type III Nrg1 heterozygous mice also show other endophenotypes of schizophrenia such as pre-pulse inhibition deficits, increased ventricle volume, altered hippocampal activity and connectivity (Wolpowitz, Mason et al. 2000, Chen, Hancock et al. 2010, Nason, Adhikari et al. 2011) indicating a possible connection between developmental Nrg1 back signaling and cellular and behavioral endophenotypes of severe neurodevelopmental disorders such as schizophrenia.

A missense mutation in *Nrg1* resulting in a single amino acid substitution in a transmembrane valine (to leucine; rs74942016-results in Val Leu at position 321 in Type III Nrg1 β1a) was found to be associated with schizophrenia (and more strongly with psychosis) in a Costa Rican population (Walss-Bass, Liu et al. 2006). Intriguingly, gamma secretase cleaves between cysteine 320 and valine 321, and leucine substitution for V_321_ results in impaired cleavage and reduced nuclear translocation of the ICD (Chen, Hancock et al. 2010, Fleck, Voss et al. 2016). This presents the intriguing possibility that loss of a specific signaling modality within this multifunctional protein might contribute to some features of neurodevelopmental disorders such as schizophrenia.

To isolate the effects of nuclear back-signaling by the Nrg1 ICD on neuronal development and transcriptomic regulation we generated transgenic mice harboring the V_321_L mutation in Nrg1. The dentate gyrus (DG) serves as the gate for incoming information into the hippocampus. Neuronal development is an ongoing process in the DG, owing to the presence of a neurogenic niche in the postnatal brain, thereby allowing us to broadly sample effects on various stages of neuronal development from neuronal fate commitment to maturation. Thus, we chose the DG as a candidate region to begin investigating the effects of impaired gamma-secretase processing of Nrg1.

## RESULTS

### Generation of the Nrg1 V_321_L mutant mouse to study γ-secretase-mediated Nrg1 signaling

Nrg1 proteins can back signal through at least two different mechanisms – 1. Local activation of PI3K-Akt signaling and 2. Nuclear translocation of a free C-terminal fragment generated by gamma secretase cleavage. The V_321_L mutation disrupts the preferred gamma secretase cleavage site (**Figure 1A**). To examine the effect of the V_321_L substitution on Nrg1 signaling, we generated a knock-in germline mutation in mice. The V_321_L mutation (gtg ttg) was introduced into C57Bl6 embryonic stem cells using a bacterial artificial chromosome (BAC) construct (**Figure 1B-D**). Heterozygotes were interbred, and offspring were born at the expected Mendelian ratios (**Figure 1D**). Homozygous V_321_L mutants were viable and fertile, with no outward morphological or growth abnormalities. Homozygous animals were able to interbreed and yielded normal-sized litters with offspring that appeared healthy. The V_321_L mutation was also viable on a Type III Nrg1 knock-down background, as a *Nrg1*^L/L^ x Type III *Nrg1*^+/-^ cross yielded both *Nrg1*^V/L^ and *Nrg1*^L/-^ offspring.

**Figure 1:**
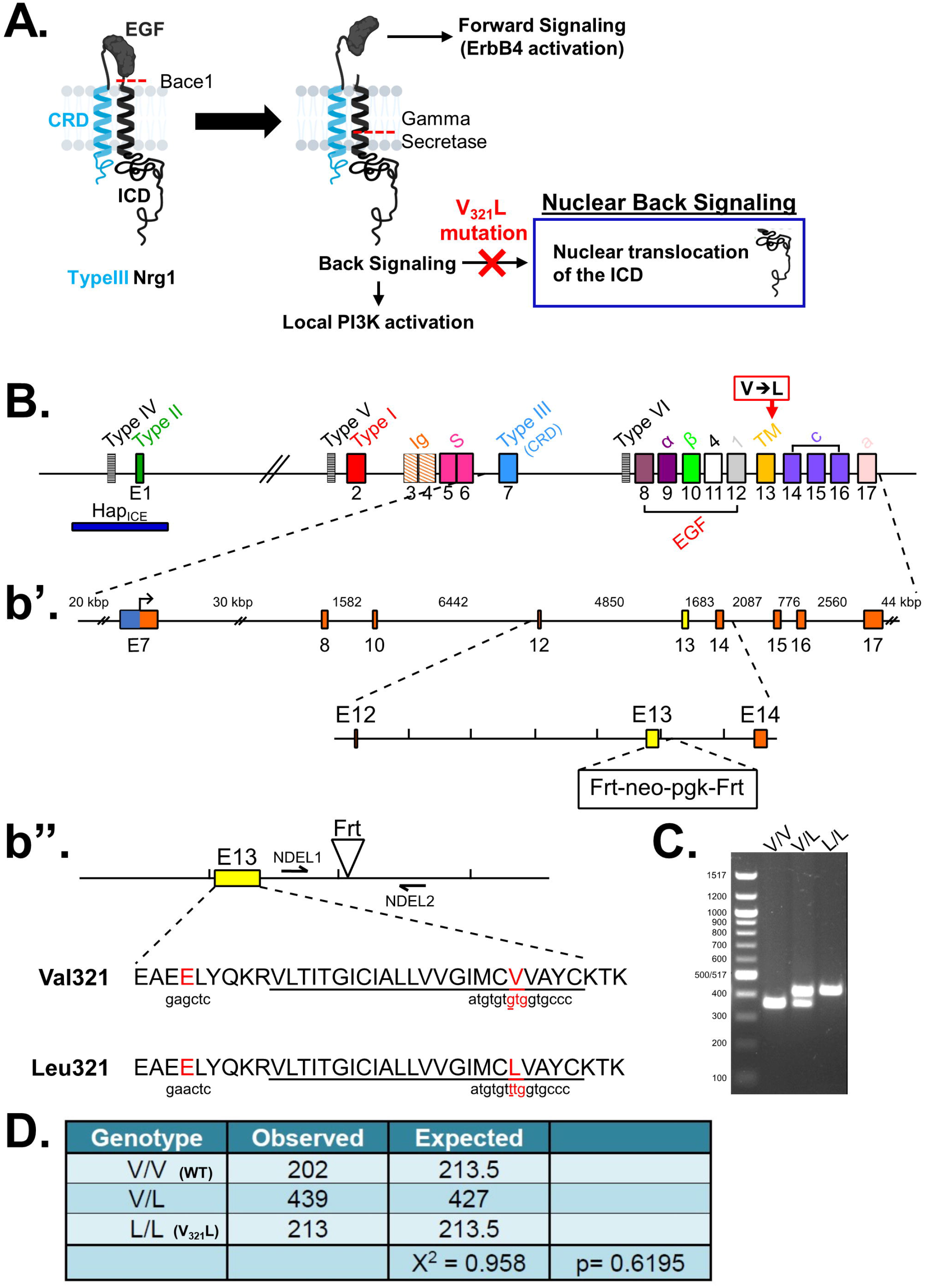
Generation of Nrg1 V_321_L knock-in mice. **(A)** A schematic of the different modes of signaling engaged by Type III Nrg1. Cleavage by Bace1 is an essential step to generate a substrate for gamma secretase. The extracellular EGF-like domain of Nrg1 can interact with ErbB4 on neighboring cells to engage the canonical forward signaling by activating ErbB4. ErbB4 interaction can also result in activation of back signaling by Nrg1. Local back signaling engages PI3K signaling at the membrane, whereas cleavage by gamma secretase results in liberation of the C-terminal ICD, which can translocate to the nucleus-nuclear back signaling. The V_321_L mutation blocks cleavage by gamma secretase and is predicted to disrupt nuclear back signaling. **(B)** A schematic diagram of the *Nrg1* genomic structure. The *Nrg1* gene encodes six families of isoforms as a result of alternative promoter usage. Types I, II, IV and V contain an Ig-like domains (encoded by exons 3 & 4). Exon 7 is the unique 5-prime coding exon for Type III Nrg1, encoding an N-terminal, cysteine-rich transmembrane domain. Exon 8, in combination with exon 9 or 10, and various combinations of exons 11, 12, and 13, encodes an EGF-like domain common to all Nrg1 isoforms. Exon 13 encodes a common C-terminal transmembrane domain. The C-terminal intracellular domain is encoded by exons 14-17. The missense SNP that results in a valine-to-leucine substitution in the C-terminal transmembrane domain Is shown above the gene. **(b’)** Schematic diagram of the BAC clone used for generating the target construct corresponding to a region of the Nrg1 gene that comprises the entire Type III coding region. Below the BAC clone is a diagram of the targeting construct, including a 5.8-kbp left homology arm, a neoR cassette in the antisense orientation flanked by Frt sites, and a 2.5-kbp right homology arm. **(b’’)** Diagram of wildtype and mutant alleles (the mutant allele following flippase removal of the neo cassette). The C-terminal transmembrane domain sequence is underlined. Black arrows labeled “NDEL1” and “NDEL2” indicate approximate genomic locations of genotyping primers. **(C)** An example genotyping gel of offspring from a het x het cross illustrating a wildtype, heterozygote and homozygote. **(D)** The mutant mouse line was maintained by breeding heterozygotes. The genotype of pups from >100 litters was analyzed for deviation from the expected 1:2:1 ratio. No deviation was found. Mice with the genotype V/V are referred to as WT in the manuscript and L/L are referred to as V_321_L as denoted in parenthesis.

### Neurons from Nrg1 V_321_L mutant mice show diminished nuclear back signaling and lack of dendritic growth in response to stimulation with soluble ErbB4

The V_321_L mutation in Nrg1 impairs gamma-secretase mediated cleavage in vitro (Fleck, Voss et al. 2016). We performed subcellular fractionation to isolate nuclei and membrane fractions from cortical and hippocampal homogenates of V_321_L homozygous mutant and their wildtype (WT) litter- and cage-mates and compared the levels of Nrg1 ICD by immunoblot analysis (**Figures 2A, 2B**). Nuclei isolated from V_321_L animals showed lower levels of Nrg1 ICD compared to nuclei from wildtype counterparts (**Figure 2A**). Concurrently, we found higher levels of full length (FL) and the membrane bound ICD (TM-ICD) in the membrane fraction of V_321_L mice compared to WT mice (**Figure 2B**). These findings indicate diminished Nrg1 nuclear back signaling in the mutant cortex and hippocampus. We next assessed stimulus-induced nuclear back signaling by addition of soluble ectodomain of the Nrg1 receptor, Erbb4 (sB4), to cultured hippocampal neurons.

**Figure 2:**
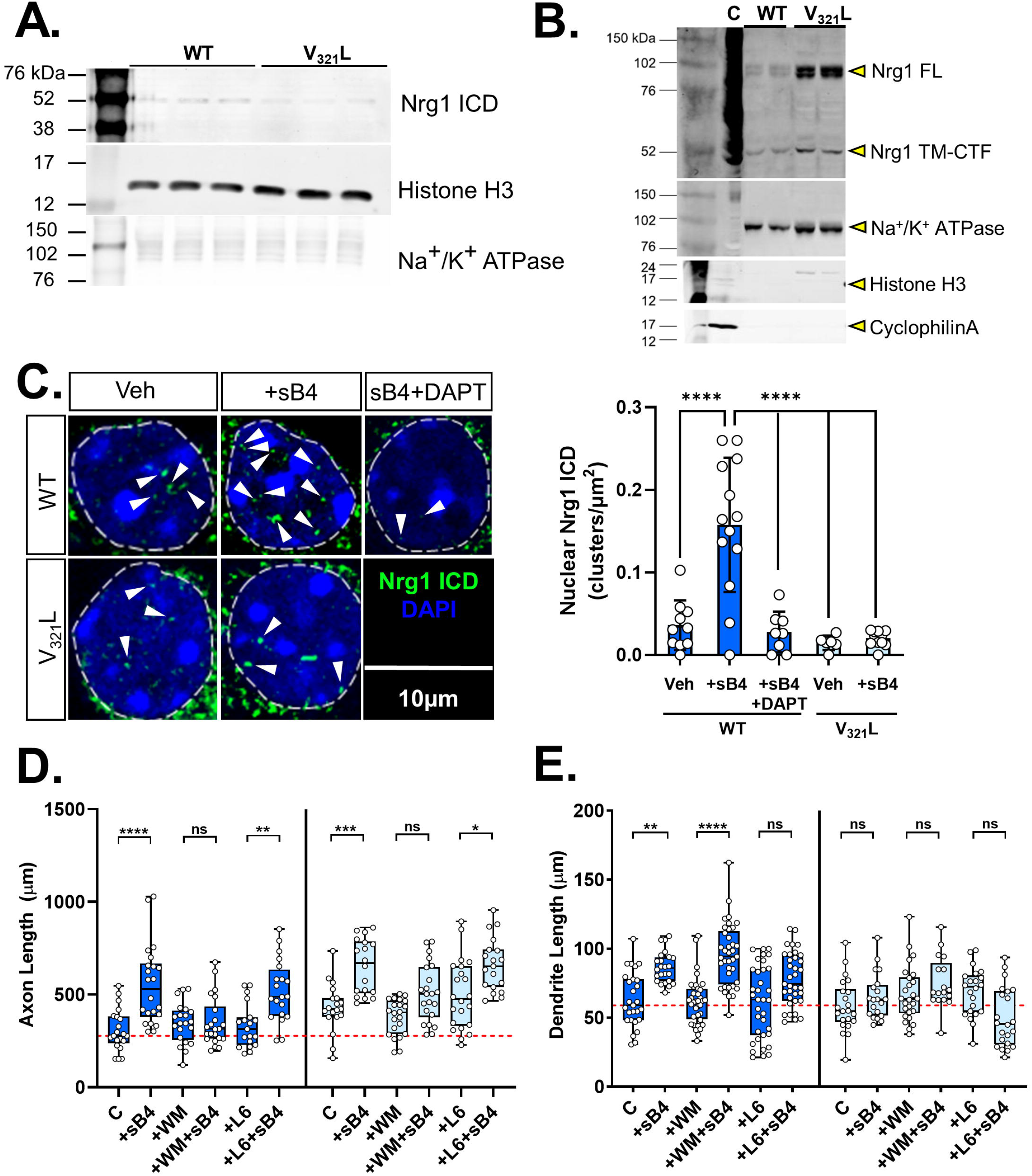
The V_321_L substitution decreases nuclear back-signaling. **(A)** Immunoblot of triplicate nuclear fractions isolated from pooled cortical and hippocampal lysates. Nrg1 ICD was detected using Santa Cruz Biotechnology antibody sc-348. Histone H3 served as a nuclear loading control, and Na+/K+ ATPase as a marker for the membrane fraction (N=3 mice/genotype). **(B)** Immunoblot of two replicates of membrane fractions isolated from pooled cortical and hippocampal lysates. NRG1 ICD was detected using Santa Cruz Biotechnology antibody sc-348. Na+/K+ ATPase served as a marker for the membrane fraction; note the lack of the nuclear marker Histone H3 or cytoplasmic marker CyclophilinA indicating clean membrane preps. Full length Nrg1 is indicated with a yellow arrowhead as “Nrg1-FL” and the membrane bound C-terminal fragment not cleaved by gamma secretase is indicated as “Nrg1 TM-CTF”. A positive control consisting of total lysate from N2A cells transfected with a Type III Nrg1 plasmid is shown in the lane labeled “C”. (N=2 mice/genotype). **(C) (Left)** Hippocampal neurons from WT (dark blue) and V_321_L (light blue) neonatal pups (P4) were cultured for 17 days in vitro and were stimulated with either vehicle (Veh), 20nM sERBB4 (sB4), or 20nM sErbB4 after a 24h pre-treatment with 20μM of the gamma secretase inhibitor DAPT (DAPT). Neurons were fixed and stained using an antibody directed against the Nrg1-ICD and counterstained with DAPI. Scale bar=10μm. **(Right)** Quantification of nuclear clusters of Nrg1-ICD. Neurons from WT mice show increased nuclear ICD clusters in response to sB4 stimulation, which is counteracted by pretreatment with DAPT (DAPT). Neurons from V_321_L mice do not respond to sB4 stimulation (N= 6-13 neurons, 3 platings/mouse, 3 mice/genotype. One-way ANOVA p-values corrected for multiple comparisons using Tukey’s posthoc test; WT Veh vs. WT sB4 p<0.0001 (****); WT sB4 vs. WT sB4+DAPT p<0.0001 (****); WT sB4 vs. V_321_L Veh p<0.0001 (****); WT sB4 vs. V_321_L B4 p<0.0001 (****). All other comparisons are statistically not statistically significant. **(D)** Cortical neurons from embryonic WT (dark blue) and V_321_L mice (light blue) (E18.5) were cultured for 3 days *in vitro* and were stimulated with soluble ErbB4 (sB4), PI3K inhibitor Wortmannin (WM), gamma secretase inhibitor L-685,458 (L6), WM+B4, or L6+B4. Neurons that underwent no drug treatments/sB4 stimulation are indicated as the control group (C). Neurons were fixed and axonal length was quantified. (Two-way ANOVA w/Tukey’s post hoc correction: WT C v. WT B4 p=0.0002 (****); WT L6 v. WT L6+B4 p=0.0047 (**); V_321_L C v. V_321_L B4 p=0.001 (***), V_321_L WM v. V_321_L WM+B4 p=0.1, V_321_L L6 v. V_321_L L6+B4 p=0.03 (*)). N=20-37 neurons per genotype per condition. ns=not significant. **(E)** Treatment and conditions same as in **D**., quantification is for dendritic length. (Two-way ANOVA w/Tukey’s post hoc correction: WT C v. WT B4 p=0.002 (**); WT WM v. WT WM+B4 p<0.0001(****)). N=20-37 neurons per genotype per condition. ns=not significant.

Type III Nrg1 back-signaling results in the appearance of distinct clusters of the ICD in the nucleus (Bao, Wolpowitz et al. 2003). To determine whether nuclear ICD clusters were altered in neurons from V_321_L mice in response to ErbB4, we cultured dispersed hippocampal neurons from postnatal day 4 (PND 4) WT and V_321_L mice (culture age:17 days *in vitro* (DIV)). We stimulated these cultures with soluble recombinant ectodomain of ErbB4 (sB4) and quantified nuclear ICD clusters (**Figure 2C left**). We found that stimulation with sB4 increased the number of nuclear ICD clusters in neurons from WT mice (**Figure 2C right;** WT Veh v. WT sB4 One-way ANOVA Bonferroni adj. p<0.0001). This increase in nuclear ICD clusters was blocked by pre-treatment with the γ-secretase inhibitor, DAPT (**Figure 2C right;** WT Veh v. WT (sB4) + DAPT Bonferroni adj. p>0.9999). We did not observe increased ICD clusters in neurons from V_321_L mice treated with sB4 (**Figure 2C right;** V_321_L Veh v. V_321_L (sB4) Bonferroni adj. p>0.9999).

As mentioned before, Type III Nrg1 back-signaling operates via two known mechanisms – 1. Local axonal, PI3K-Akt signaling and 2. γ-secretase-dependent nuclear signaling, required for dendritic growth and complexity (Chen, Hancock et al. 2010, Fazzari, Snellinx et al. 2014). We asked whether neurons from V_321_L mice are selectively deficient in dendrite development in response to ErbB4 stimulation. Cortical neurons from embryonic (E18.5) WT and V_321_L mouse pups were cultured for 3 days *in vitro*. On the third day, neurons were treated with sB4 with or without pharmacological inhibition of PI3K using wortmannin (WM) or γ-secretase using L-685,458 (L6).

Neurons from WT mice showed increases in axonal and dendritic length in response to ErbB4 treatment which were blocked by WM and L6 respectively (**Figure 2D and 2E dark blue boxes**; axonal length: WT Control v. WT sB4 p=0.0002; dendritic length: WT Control v. WT sB4 p=0.0019). WM treatment did not block ErbB4-induced dendritic growth and L6 treatment did not prevent ErbB4-induced axonal growth (axonal length: WT L6 v. WT L6+sB4 p<0.0001; dendritic length: WT WM v. WT WM+sB4 p<0.0001). These results agree with previously published data showing that Nrg1 nuclear back-signaling influences dendritic growth (Chen, Hancock et al. 2010). These data also establish that stimulation of axonal growth requires PI3K signaling and that these two modes of Nrg1 back signaling are functionally independent.

Neurons from V_321_L mice showed increased axonal length in response to ErbB4 stimulation which was blocked by WM treatment and not by L6 (**Figure 2D light blue boxes**; axonal length: V_321_L Control v. V_321_L sB4 p=0.0014, V_321_L WM v. V_321_L WM+sB4 p=0.1113, V_321_L L6 v. V_321_L L6+sB4 p=0.0289). On the other hand, V_321_L neurons did not show increases in dendritic length in response to ErbB4 stimulation (**Figure 2E** dendrite length: V_321_L Control v. V_321_L sB4 p>0.9999). These results indicated that PI3K signaling-dependent axonal growth was intact in V_321_L mutant mice with a selective disruption of γ-secretase-dependent dendritic growth.

Thus, V_321_L mice show disruptions to regulated intramembrane proteolysis of Nrg1 and thereby to nuclear translocation of the ICD. Additionally, we demonstrate that dendritic growth in response to sErbB4 stimulation was absent in neurons from V_321_L mutant mice whereas axonal growth in response to sErbB4 was intact.

### Developmental regulation of Nrg1 nuclear back signaling in granule cell cultures

In hippocampal neurons, the Type III Nrg1 protein is part of the presynaptic membrane where it is predicted to interact with ErbB4 on dendrites of GABAergic interneurons (Vullhorst, Neddens et al. 2009, Vullhorst, Ahmad et al. 2017). We noted the presence of nuclear Nrg1 ICD clusters at baseline in our previous experiments, which increased in number following stimulation with ErbB4 (**Figure 2C**). Thus, we next sought to examine this baseline endogenous nuclear signaling and asked whether it might correspond to a specific developmental window. We first characterized our P4 hippocampal culture preparation to ask if GABAergic interneurons were present in our culture, as a possible source of ErbB4 in the culture. ∼60-70% (Mean: 63% ± 10% std.dev) of the neurons were GCs (Prox1+) whereas ∼25% (Mean: 25% ± 7.5% std.dev) of the neurons were GABAergic (GAD67+) and the remainder ∼12% (Mean: 12%± 12% std.dev) were other neurons likely corresponding to glutamatergic pyramidal and mossy cells (**Figure 3A**). Next, we assessed baseline and stimulated nuclear back signaling at DIV 10, 14, and 17. We found that the baseline nuclear back signaling diminished over time in culture but remained inducible by sErbB4 treatment (**Figure 3B**; Kruskal-Wallis Test p= 0.0003 KW= 23.31; DIV10 Control vs. DIV17 Control Dunn’s adj.p=0.049, DIV17 Control vs. DIV17 +sB4 Dunn’s adj.p=0.006, DIV10 +sB4 vs. DIV17 +sB4 Dunn’s adj.p>0.9999). Intriguingly, at DIV10 the level of baseline nuclear ICD clusters was high and sErbB4 treatment could not significantly enhance this signal (Dunn’s adj.p>0.9999). Since Nrg1 nuclear back signaling is thought to be regulated by cell-cell interactions, and in turn influences neurite growth and development, we characterized the axonal and dendritic growth over time in our GC cultures to ask whether the high basal nuclear back signaling might correspond to a particular developmental process associated with neurite growth. Axonal growth (indicated by the area covered by SMI312+ processes) increased from DIV1 to DIV10, plateauing thereafter (**Figure3C**; Kruskal-Wallis p=0.0001, KW=25.79; DIV1 vs. DIV10 Dunn’s adj.p=0.0003; DIV1 vs. DIV14 Dunn’s adj.p<0.0001; DIV5 vs. DIV14 Dunn’s adj.p=0.0462). Qualitatively, we noted a gradual increase in axonal bundling over time. Similarly, we noted a gradual increase in dendritic growth (indicated by the area covered by MAP2+ processes) from DIV1 to DIV14 (**Figure 3D**; Kruskal-Wallis p>0.0001, KW=24.93; DIV1 vs. DIV10 Dunn’s adj.p=0.0035; DIV1 vs. DIV14 Dunn’s adj.p<0.0001; DIV5 vs. DIV14 Dunn’s adj.p=0.0308). Qualitatively, we also noted an enhancement of dendritic complexity between DIV10 and 14 (**Figure 3D**). As noted earlier, nuclear back signaling was maximal at DIV10, thus we examined interactions between axons and dendrites at DIV5,10 and 14 to assess whether an increase in axon-dendrite contacts corresponds to the high nuclear back signaling (**Figure 3E**). We found that at DIV5 a few thin SMI312+ processes (individual axons) were in proximity to MAP2+ processes (dendrites) (**Figure 3E, left**). This was dramatically enhanced at DIV10 where we noted a higher amount of axonal coverage, along with multiple axons “running along” dendrites (**Figure 3E, middle**). Finally, at DIV14 we noted axonal bundling indicated by the presence of thicker SMI312+ processes, which ran along dendrites (**Figure 3E, right**). Thus, under these culture conditions, DIV10 represents a period of dynamic axonal growth, axon-dendrite contact, and endogenous nuclear back signaling, which is followed by increased dendritic complexity. Intriguingly, this period falls squarely within the gamma secretase inhibition sensitive window for dendrite development (see below) indicating that the nuclear ICD might regulate genes related to dendrite development.

**Figure 3:**
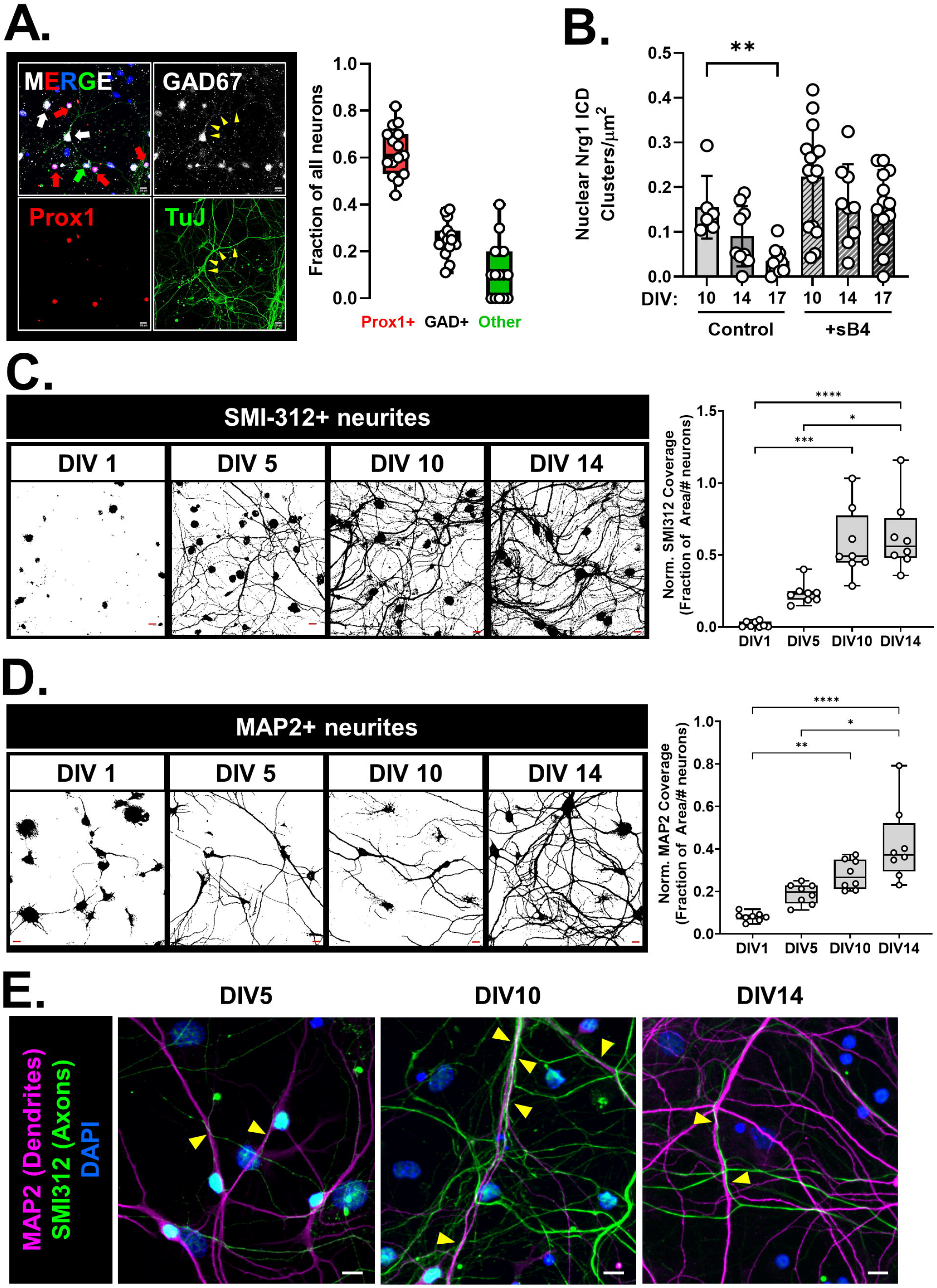
The peak of Nrg1 nuclear back signaling aligns with the peak of axon growth and precedes the peak of dendrite growth. **(A)** Characterization of neuronal types present in the P4-5 hippocampal culture. (**Left**) Representative image from a WT P5 culture stained for GAD67 (greyscale; GABAergic neuron marker), TuJ (green; neuronal marker), Prox1 (red; granule cell (GC) marker), and counterstained with DAPI (blue; to label nuclei). GCs were identified as Prox1+ TuJ+ cells. GABAergic interneurons were identified as GAD67+ TuJ+ Prox1-cells, where the GAD67 staining also delineated the neurites (example indicated by yellow arrowheads). Neurons not stained by Prox1 or GAD67 were identified as other excitatory neurons likely representing pyramidal and/or mossy cells. Scale bar = 10µm. (**Right**) Quantification of neuronal types as proportion of all neurons. Data were pooled from cultures imaged on various days in vitro (DIV 5,10,14). Overall, GCs make up majority of the neurons in these cultures (mean: 62.6% std.dev: 10.5%). N=15-16 platings from 12-16 mice from 2 litters. **(B)** Quantification of Nrg1 ICD clusters in nuclei of WT neurons from P4-5 hippocampal cultures at different DIVs under basal conditions (Control) and after stimulation with sB4 (20nM). N=6-15 neurons/ condition from platings made from 3 mice; Kruskal-Wallis test p=0.005 (**) KW=10.59. (Dunn’s corrected multiple comparisons) DIV10 vs. DIV17 p=0.0047. <u>Note</u>: DIV17 group same as Figure 2C for comparison. **(C)** (**Left**) Representative thresholded images of SMI-312 staining (pan-axonal neurofilament marker) from WT mice at DIV 1, 5, 10, and 14. Scale bar (red) = 10µm. (**Right**) Quantification of fraction of area covered by SMI-312 staining normalized to the number of neurons in the imaging field. In WT cultures overall axonal coverage increased until DIV10 stabilizing thereafter. Kruskal-Wallis test p<0.0001 KW=25.79. (Dunn’s corrected multiple comparisons) DIV1 vs. DIV10 p=0.0003 (***), DIV1 vs. DIV14 p<0.0001 (****), DIV5 vs. DIV14 p=0.046 (*). All other comparisons were not significant. N=8-10 platings from 6-8 mice from 2 litters for each time point. **(D)** (**Left**) Representative thresholded images of MAP2 staining (dendrite marker at time points after DIV1) from WT mice at DIV 1, 5, 10, and 14. Scale bar (red) = 10µm. (**Right**) Quantification of fraction of area covered by MAP2 staining normalized to the number of neurons in the imaging field. Overall dendrite coverage increased till DIV14 in WT cultures. Kruskal-Wallis test p<0.0001 KW=24.93. (Dunn’s corrected multiple comparisons) DIV1 vs. DIV10 p=0.004 (**), DIV1 vs. DIV14 p<0.0001 (****), DIV5 vs. DIV14 p=0.03 (*). All other comparisons were not significant. N=8-10 platings from 6-8 mice from 2 litters per genotype for each time point. **(E)** Representative images of MAP2 and SMI312 staining of WT cultures at DIV5, 10, and 14 showing increased axon-dendrite interactions at DIV10. Yellow arrowheads denote regions of axon-dendrite contacts. Scale bar= 10µm.

### Gamma secretase activity antagonistically regulates axon and dendrite development

The V_321_L mutation disrupts gamma secretase processing of Nrg1 thereby preventing nuclear translocation of the ICD (**Figure 2C**). Additionally, nuclear translocation of the ICD is regulated during development in GC-enriched cultures (**Figure 3B**). Thus, we next queried the role of gamma secretase activity in axon and dendrite development during specific developmental periods. We noted that the levels of Nrg1 ICD in the nucleus peak at or prior to DIV10 after which they decline (**Figure 3B**). Additionally, it was after DIV10 that we noted the dramatic increase in dendritic complexity (**Figure 3D, left**). Thus, we hypothesized that inhibiting gamma secretase during this period might influence dendrite development. We treated P4 WT hippocampal cultures with the gamma secretase inhibitor, DAPT, or vehicle (control) for varying durations of time – DIV1 to 14, DIV5 to 14, or DIV12 to 14 (**Figure 4A, left**). Cultures were fixed and stained for the pan-axonal marker SMI312 and dendritic marker MAP2 on DIV14 to quantify axonal and dendritic coverage (**Figure 4A, right**). Inhibiting gamma secretase prior to DIV12 resulted in severe dendritic growth defects (**Figure 4B, top; Figure 4C**, Kruskal-Wallis test p=0.0003 KW=19.09; Control vs. DAPT1-14 p=0.015; Control vs. DAPT5-14 p=0.003; Control vs. DAPT12-14 p>0.9999; DAPT5-14 vs. DAPT12-14 p=0.011). Starting gamma secretase inhibition after DIV5 was sufficient to induce the severe dendritic growth defect phenotype, indicating that there is a period of gamma secretase-dependent dendrite growth between DIV 5 and 12. We also found that there was an increase in the number of primary neurites immunoreactive for the pan-axonal marker, but no significant differences in axonal coverage between the control and gamma secretase treated groups (**Figure 4B, bottom; Figure 4D**: # SMI312+ primary neurites, Kruskal-Wallis test p=0.01, KW=11.13. C vs. DAPT5-14 p=0.04; **Figure 4E**: Axonal coverage, Kruskal-Wallis test p=0.21 KW=4.487). These results indicate that gamma secretase activity between DIV5 and 12 is necessary to constrain axonal development and promote dendritic development.

**Figure 4:**
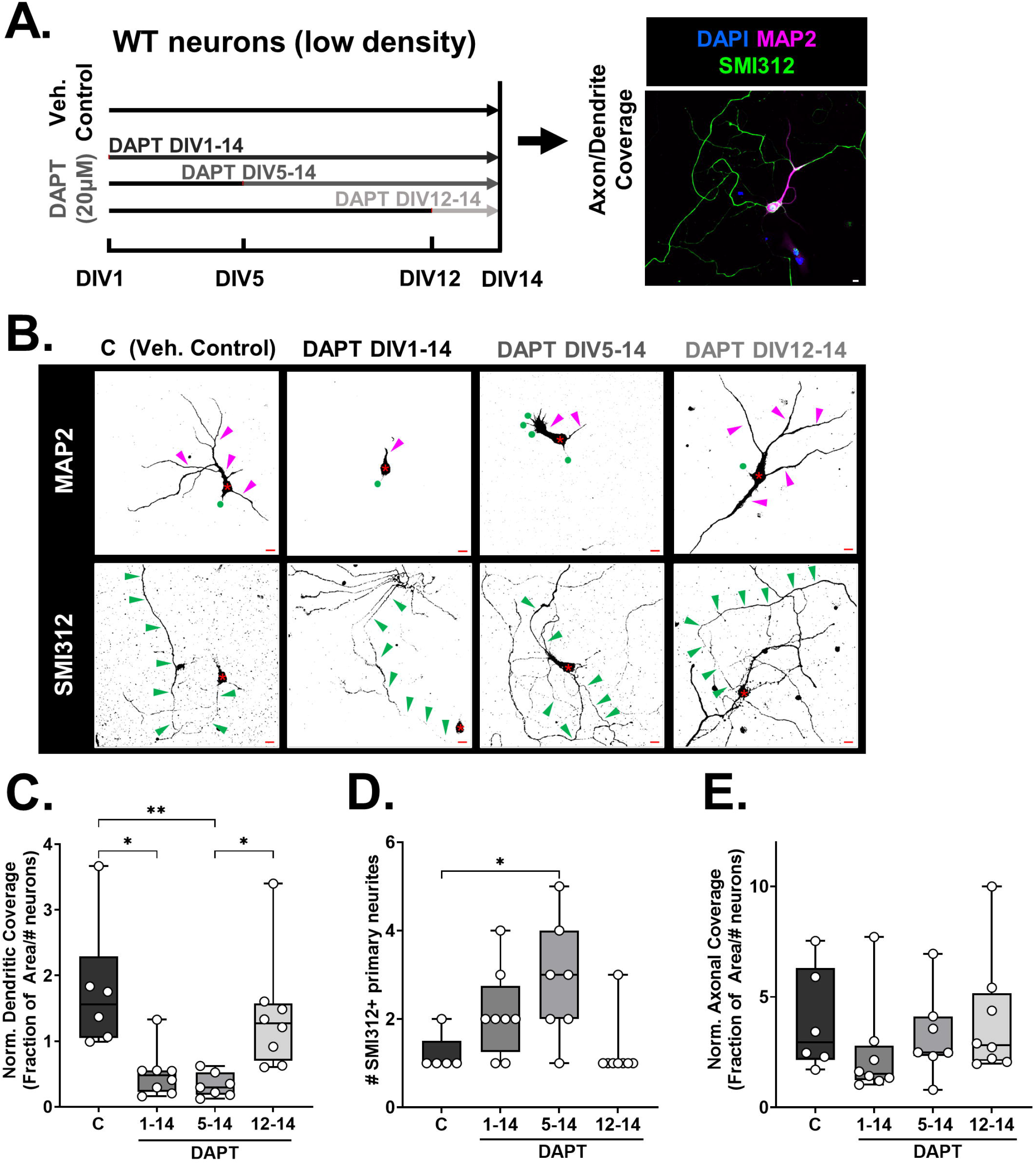
Gamma secretase regulates axon and dendrite development. **(A)** Schematic of DAPT treatment regimens that cultured neurons from WT mice (P4/5 hippocampus) were subjected to. Vehicle controls (C) received equal volume of DMSO diluted in culture media. DAPT1-14 group received DAPT treatment from DIV 1-14, DAPT5-14 group was cultured for the first 5 days without DAPT but received DAPT from DIV5 onwards, and DIV12-14 group received DAPT from DIV12 onwards. All platings were collected at DIV14, fixed and stained for MAP2 and SMI312. **(B)** Representative images of **(Top)** MAP2 and **(Bottom)** SMI312 immunostaining for each of the treatment conditions described in **panel A**. Asterisk (*) denotes the soma of each neuron. Magenta and green arrow heads indicate MAP2+ and SMI312+ neurites respectively. Green circles in the MAP2 stained images mark the point(s) of origin of SMI312+ neurites. The same neuron is shown in both Top and Bottom panels. Note that for the DAPT DIV1-14 images, the SMI312 panel was recentered to allow visualization of the axon (soma now in bottom right corner). Scale bar= 10µm. **(C)** Quantification of fraction of area covered by MAP2 staining normalized to the number of neurons in the imaging field. Inhibition of gamma secretase prior to DIV12 significantly reduced dendritic growth. Kruskal-Wallis test p=0.0003 KW=19.09. C vs. DAPT1-14 Dunn’s corrected p=0.015 (*), C vs. DAPT5-14 Dunn’s corrected p=0.003 (**), DAPT5-14 vs. DAPT12-14 Dunn’s corrected p=0.011 (*) N=6-8 platings from 6-8 mice from 2 litters per genotype for each time point. **(D)** Quantification of number of primary neurites immunoreactive for SMI312 after vehicle control treatment or treatment with DAPT for varying durations. Kruskal-Wallis test p=0.01, KW=11.13. C vs. DAPT5-14 Dunn’s corrected p=0.04 (*). All other comparisons were noy statistically significant. N=6-8 platings from 6-8 mice from 2 litters per genotype for each time point. **(E)** Quantification of fraction of area covered by SMI312 staining normalized to the number of neurons in the imaging field. Inhibition of gamma secretase had no significant effect on axonal coverage. Kruskal-Wallis test p=0.21 KW=4.487. N=6-8 platings from 6-8 mice from 2 litters per genotype for each time point.

### Changes in Nrg1 V_321_L mutant dentate gyrus transcriptome point to aberrant neurogenesis, cell cycle dynamics, and dendrite development

The Nrg1 ICD has strong transactivation properties (Bao, Wolpowitz et al. 2003) and V_321_L mice have impaired nuclear translocation of the ICD (**Figure 2**). Thus, we predicted that impaired nuclear signaling by the Nrg1 ICD would result in transcriptomic changes. We extracted RNA from microdissected DG from WT and V_321_L mice for RNA sequencing (**Figure 5A**). Differential gene expression analysis revealed 1312 significantly dysregulated genes (colored dots, ANCOVA corrected p≤0.1 and linear fold change of ±1.25) between WT and V_321_L mice among which were genes specifically important for DG granule cell specification and function such as *Prox1*, *Calb1*, and *Synpr* (**Figure 5B**; **Table 5-1**). To identify possible functional alterations that might result from transcriptional dysregulation in V_321_L mice, we used ingenuity pathway analysis (IPA) to find cellular processes that might be affected by the significantly dysregulated genes. Several processes along the neurodevelopmental trajectory were predicted to be disrupted in the V_321_L DG, including cell proliferation, differentiation, and dendrite development (**Table 5-2**). Disease annotations in IPA indicated that dysregulated genes in the V_321_L DG were enriched in genes associated with schizophrenia-susceptibility, indicating that the V_321_L mutation alters the transcriptional landscape in ways that have been associated with the genetic architecture of schizophrenia (**Table 5-3**). We also found that dysregulated genes in V_321_L mice were enriched in cancer-associated genes, consistent with predicted changes in regulation of cell proliferation (**Table 5-3**).

**Figure 5:**
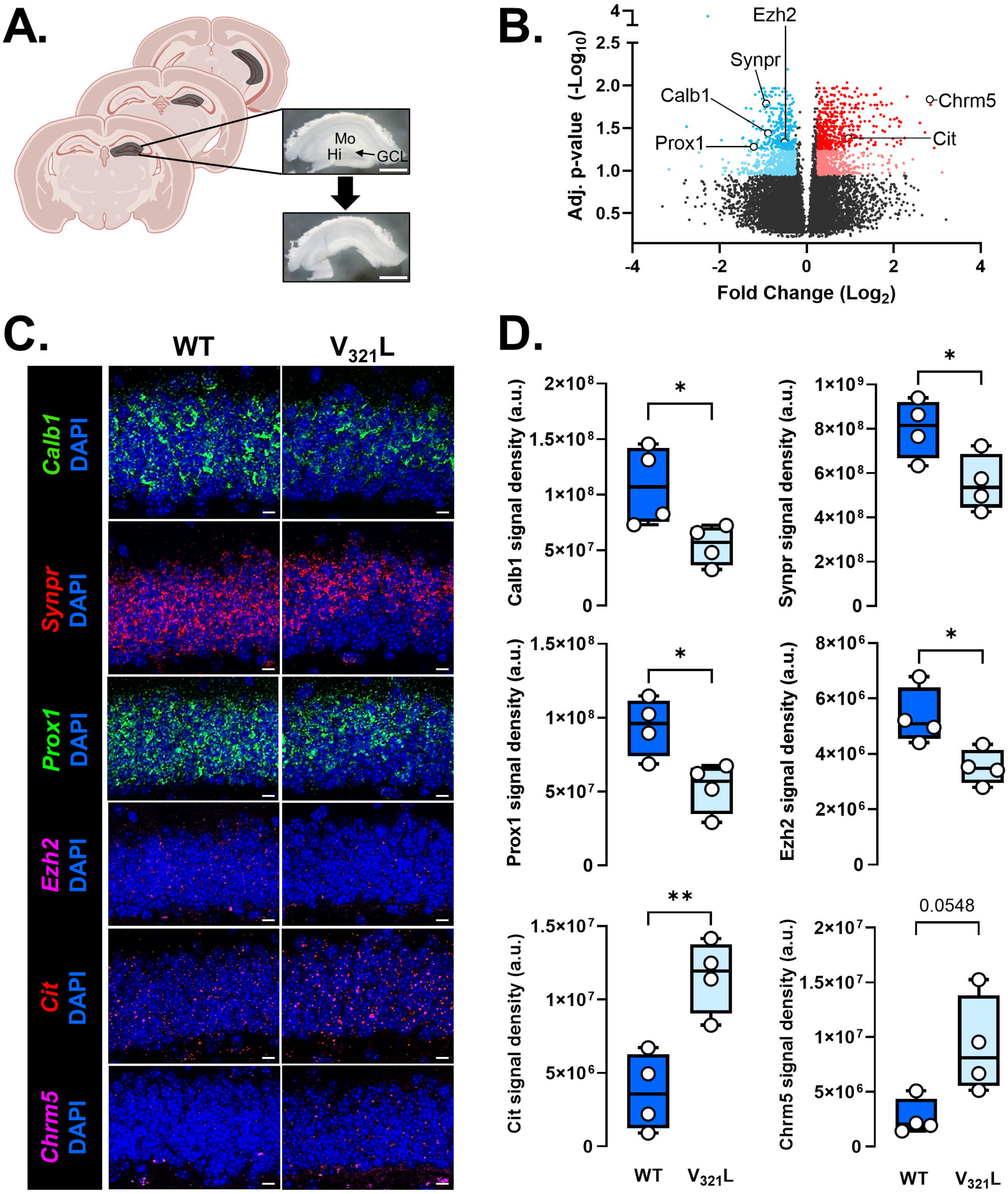
The V_321_L substitution in Nrg1 alters gene expression in the DG. **(A)** Schematic and representative image of DG dissection from hippocampal slices; Mo= molecular layer, GCL= granule cell layer, Hi= hilus. **(B)** Volcano plot for differentially expressed genes between V_321_L and WT DG. Each datapoint represents a single gene. Colored points delineate fold change of at least 1.25 (blue dots – downregulated, red dots-upregulated in the mutant samples) with an adjusted p-value <0.1, and dark colored points represent those with ANCOVA adjusted p-value <0.05. (N=6 mice/genotype. 3 Male/3 Female). *Prox1*, *Calb1, Synpr, Ezh2, Cit,* and *Chrm5* were chosen for validation via RNAScope *in situ* hybridization (ISH). The complete list of differentially expressed genes, statistics, and pathway analyses can be found in **Extended Data Tables 5-1, 5-2, 5-3**. **(C)** Representative images of WT and V_321_L sections showing DAPI stained DG granule cell layer (GCL) and staining for mRNA of the indicated genes in panel B. **(D)** Quantification of RNAScope signal shown in panel A. (**From top to bottom,** WT vs. V_321_L Welch’s t-test (two-tailed)) *Calb1*: p=0.049 (*) t=2.679, df=4.435; *Synpr*: p=0.04 (*) t=2.672, df=5.992; *Prox1*: p=0.02 (*) t=3.174, df=5.882, *Ezh2*: p=0.03 (*) t=3.031, df=5.034, *Cit*: p=0.005 (**) t=4.352, df=5.983, *Chrm5*: p=0.0548 t=2.740, df=3.808. N=4 mice/genotype.

To validate the results from our RNASeq experiment we performed fluorescent in situ hybridization (FISH) using the RNAScope assay on WT and V_321_L DG brain sections and quantified the expression of a few top DEGs (indicated in **Figure 5B**) in the granule cell layer of the dorsal DG – *Prox1*, *Calb1*, *Synpr, Ezh2, Cit, and Chrm5* (**Figure 5C**). In line with our RNASeq data, expression of *Prox1*, *Calb1, Synpr, and Ezh2* was significantly lower in the V_321_L DG compared to WT (**Figure 5D**, *Prox1*: WT v. V_321_L p=0.02, t=3.174, df=5.882; *Calb1*: WT v. V_321_L p=0.05, t=2.679, df=4.435; *Synpr*: WT v. V_321_L p=0.04, t=2.672, df=5.992; *Ezh2*: WT v. V_321_L p=0.03, t=3.031, df=5.034) and expression of *Cit* and *Chrm5* was higher (**Figure 5D**, *Cit*: WT v. V_321_L p=0.005, t=4.352, df=5.983; *Chrm5*: WT v. V_321_L p=0.05, t=2.740, df=3.808). We did not find any significant differences in the DAPI+ GCL area between WT and V_321_L mice indicating no overt difference in numbers of GCs between the genotypes (p=0.3429 Mann-Whitney Rank Sum Test, U=4).

### Shared transcriptional regulatory mechanisms among DEGs in the V_321_L DG implicate dysregulation of the polycomb repressor complex 2 (PRC2)

We next asked whether the differentially expressed genes shared transcriptional regulatory mechanisms using ENCODE and ChEA databases (Lachmann, Xu et al. 2010, Chen, Tan et al. 2013, Kuleshov, Jones et al. 2016, Xie, Bailey et al. 2021). The 664 significantly upregulated genes in V_321_L mice were enriched for genes predicted to be regulated by members of the polycomb repressor complex 2 (PRC2) (SUZ12, EZH2), RE1-Silencing Transcription factor (REST), and the CTCF-cohesin complex (CTCF, RAD21, SMC3) involved in chromatin looping (**Table 5-4**). We grouped these regulators into two higher order groups: 1. PRC2+REST and 2. CTCF-Cohesin. We found that 126 of the 664 upregulated genes were targets for regulation by the PRC2 and 81 of the 664 upregulated genes were targets of REST; 28 of these 81 REST-target genes (∼35%) were also targets of the PRC2. Gene ontology analyses for the PRC2-target genes revealed enrichment of genes predicted to be involved in synapse assembly, synaptic transmission, and neuronal differentiation (**Table 5-5**). Intriguingly, we found that EZH2 and EED (catalytic subunits of PRC2) were downregulated in the RNASeq comparison between WT and V_321_L DG (**Table 5-1; Figure 5D**). The CTCF-Cohesin regulated genes were enriched for functions such as axon guidance, cell migration, synapse assembly, and protein phosphorylation.

The 647 downregulated genes showed strongest enrichment of genes predicted to be regulated by the transcription factor E2F4 (odds ratio: 3), Upstream Binding Transcription Factor (UBTF), Sin3 Transcription Regulator Family Member A (SIN3A), and Myc-associated Factor X (MAX) (**Table 5-4**). The E2F4-target genes were found to be involved in mitotic spindle organization, microtubule organization, RNA-splicing, and DNA replication (**Table 5-5**). Overall, the downregulated genes were found to be involved in positive regulation of mitosis.

The observed changes in gene expression strongly implicate the Nrg1 ICD as a regulator of E2F4 and PRC2 function during a neurodevelopmental program. Based on these data we predicted that there would be altered proliferation and neural differentiation within the V_321_L mutant neurogenic niche.

### Differential gene expression in the V_321_L DG can be largely accounted for by a loss of nuclear back signaling

Bulk RNASeq of tissue offered us a snapshot into steady-state differences in gene expression between WT and V_321_L DG. To determine which of these gene expression differences, if any, might reflect a direct effect of altered nuclear back signaling, we measured the effect of stimulating nuclear back signaling on gene expression in cultured P4 hippocampal neurons at DIV10 (**Figure 6A, top**). We pre-treated the cultures with the gamma secretase inhibitor (DAPT) or vehicle on DIV9 for 24h, following which we stimulated the cultures with sErbB4 or vehicle. RNA was collected 4h after ErbB4 stimulation and sequenced. We compared the following conditions to identify Nrg1 nuclear back signaling regulated genes-1. DAPT vs. sErbB4 (overall back signaling effect adjusted for baseline signaling). 2. DAPT vs. DAPT+sErbB4 (local back signaling effect), and 3. DAPT+sErbB4 vs. sErbB4 (nuclear back signaling effect adjusted for local back signaling) (**Figure 6A bottom**). We found that local back signaling did not result in significant gene expression changes at this time point (12 DEGs). Nrg1 nuclear back signaling resulted in 663 DEGs (**Table 6-1**; **Figure 6A bottom**).

**Figure 6:**
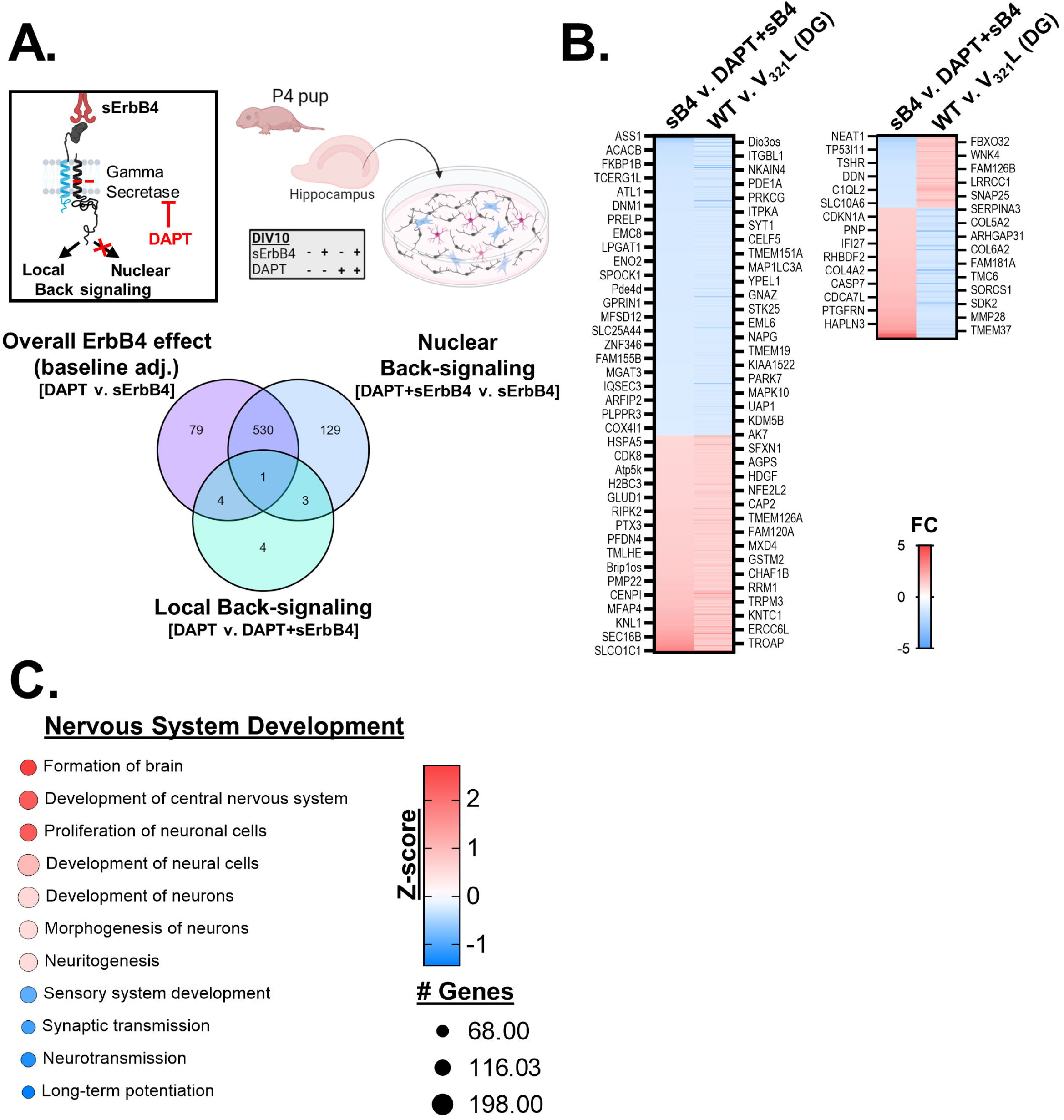
Differential gene expression in the V_321_L DG can be largely accounted for by a loss of nuclear back signaling. **(A)** (**Top**) Schematic of the experimental paradigm. P4-5 hippocampal cultures were maintained for 8-9 days after which they were treated with DMSO (vehicle control) or 20µM DAPT overnight. On DIV9-10 cultures were stimulated with 20nM soluble ErbB4 (sErbB4) or equal volume 1xPBS in culture media (vehicle control) for four hours prior to RNA extraction. The four treatment conditions are shown – Vehicle alone, sErbB4 alone, DAPT alone, DAPT+sErbB4. N=3 platings from 6-8 mice from 3 litters per genotype for each condition. (**Bottom**) Venn diagram showing overlap in the differentially expressed genes (DEGs) (fold change >1.25 in either direction, post-hoc corrected p<0.05). Local back signaling did not result in significant changes in gene expression. High overlap in the baseline adjusted ErbB4-effect and the nuclear back signaling effect indicates a high level of baseline back signaling at DIV9-10 consistent with data shown in Figure 3B. Also see **Extended Data Table 6-1, 6-2**. **(B)** (**Left**) Heatmaps showing changes in gene expression in response to sErbB4 treatment compared to expression of the same genes in WT DG tissue relative to V_321_L DG tissue for genes whose expression is changed in the same direction *in vitro* and *in vivo*. All genes detected in both datasets irrespective of the degree of fold change are displayed. Each row represents a gene, and every 10^th^ row is labeled with the gene name on either side of the heat map. The genes are ordered from most downregulated (blue) to most upregulated (red) genes from the *in vitro* experiment. Color of the heatmap represents the fold change in gene expression given by the legend displayed alongside. (**Right**) Heatmaps showing changes in gene expression in response to sErbB4 treatment compared to expression of the same genes in WT DG tissue relative to V_321_L DG tissue for genes whose expression is oppositely regulated *in vitro* compared to *in vivo*. Each row represents a gene, and every 10^th^ row is labeled with the gene name on either side of the heat map. The genes are ordered from most downregulated (blue) to most upregulated (red) genes from the *in vitro* experiment. Color of the heatmap represents the fold change (FC) in gene expression given by the legend displayed alongside. **(C)** Differentially expressed genes were subjected to pathway enrichment analysis using Ingenuity Pathway Analysis (IPA), which identified significantly enriched pathways represented by the DEGs as well as a z-score of direction in which these pathways are predicted to be altered based on the directions of changes in expression of the genes comprising each pathway. Each pathway is indicated by a circle with the size of the circle indicating the proportion of genes comprising that pathway in the IPA database that are represented in the DEG set. The color of the circles is according to the heatmaps displayed alongside indicating the adjusted p-values. Red pathways are predicted to become upregulated and blue pathways are predicted to become downregulated based on the aggregate gene expression changes effected by Nrg1 nuclear back signaling.

We next compared the change in the expression of individual genes due to the V_321_L mutation in the DG and the change in expression of the same genes from acutely stimulated nuclear back signaling (**Figure 6B**). We found that bulk of the genes found to be differentially expressed in the V_321_L DG were also acutely regulated by nuclear back signaling i.e., genes upregulated by stimulating nuclear back signaling in vitro showed higher expression in WT DG compared to the V_321_L DG and vice versa (**Figure 6B**). A smaller number of genes showed the opposite effect i.e., genes upregulated by stimulating nuclear back signaling were found to be lower in expression in the WT DG compared to V_321_L DG indicating potential compensatory mechanisms and/or other regulators of these genes (**Figure 6B**). Gene ontology analyses of the nuclear back signaling regulated genes indicated significant enrichment of genes involved in proliferation of neuronal cells and regulation of neuronal morphogenesis, consistent with predictions from the RNASeq of DG tissue from V_321_L mice (**Figure 6C**).

### Abnormal progenitor pool maintenance and differentiation in the V_321_L mutant dentate gyrus

We next quantified the effect of the V_321_L mutation on adult neurogenesis in the dentate gyrus. Stereological analysis revealed a statistically significant decrease in the proliferative population (Ki67+) in V_321_L animals compared to WT (**Figure 7A**; p= 0.03, Welch’s t-test), whereas newborn neuron (Dcx+) numbers were comparable between V_321_L and WT animals (**Figure 7B**; p=0.99, Welch’s t-test). Thus, while proliferation of stem cells was stunted in the V_321_L DG, this impairment did not result in an appreciable difference in the number of newborn neurons produced.

**Figure 7:**
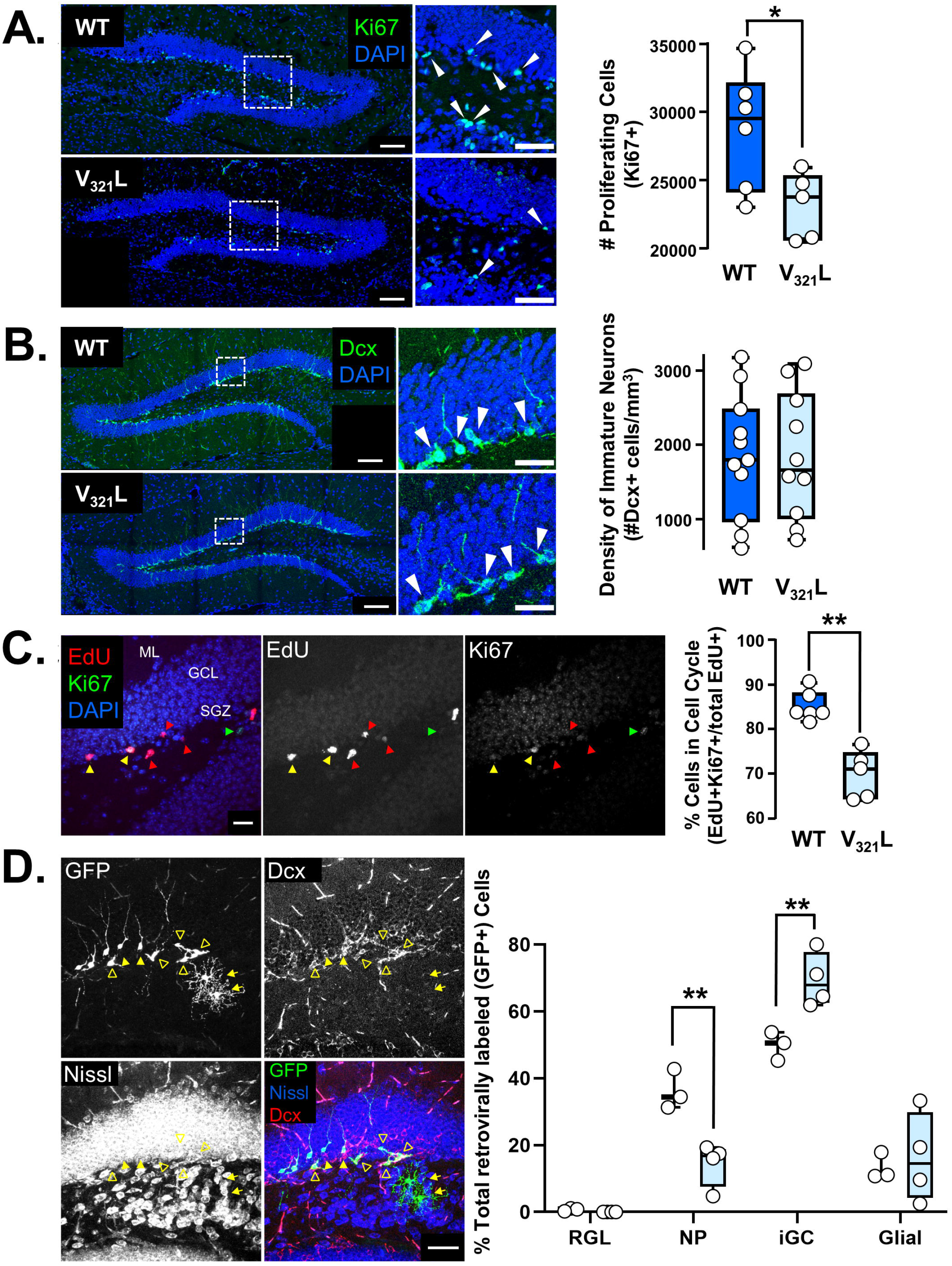
V_321_L DG shows reduced stem cell proliferation, increased cell cycle exit and commitment to neuronal fate. **(A) Left,** immunohistochemistry of Ki67 (green), a marker for proliferation, in the DG of P22 WT and V_321_L littermates and cage mates, merged with DAPI (blue) counterstain, scale bar=100μm. Inset shows a higher magnification image with arrowheads indicating Ki67+ proliferating cells in the subgranular zone of the DG, scale bar=50μm. **Right,** stereological quantification of Ki67+ populations in the DG of P21 WT and V_321_L (WT N= 6 mice, V_321_L N= 5 mice; p=0.03 (*), Welch’s t-test. t=2.697, df=8.050). **(B) Left,** doublecortin (Dcx) immunoreactivity (green) merged with DAPI (blue) counterstain in DG of WT and V_321_L mice. Inset shows higher magnification image of the region delineated in the white box; arrowheads point to Dcx+ cells in the DG. **Right,** quantification of Dcx+ cell numbers from confocal images of subsampled, coronal DG sections. Confocal images were acquired in 3-μm z-steps from 20-μm coronal sections, and 4-10 images were quantified for each animal (P60-P180 WT N= 11 mice, V_321_L =10 mice, p= 0.9865, Welch’s t-test. t=0.017, df=18.68). **(C) Left,** Image from a WT DG showing neural progenitors in S phase of the cell cycle labeled with EdU by intraperitoneal injection (50μg EdU/g body weight) in ∼P90 animals. 24 hours post-injection, samples were harvested and processed to detect Ki67 immunoreactivity. EdU and Ki67 co-immunoreactivity indicates that cells remained in the cell cycle 24 hours post-injection (yellow arrowheads). EdU immunoreactivity without Ki67 expression indicates that cells exited the cell cycle (red arrowheads). Ki67 immunoreactivity alone indicates cells that were not in S phase at the time of injection but were in the cell cycle at the time of tissue harvest. **Right,** Quantification of cell cycle reentry by computing the proportion of cells that remained in the cell cycle (EdU+ Ki67+) out of all cells in S phase of the cell cycle at the time of labeling (EdU+ cells). Epifluorescent images were acquired in 1-μm z-steps from 20-μm coronal sections, and 12-14 sections were quantified for each animal (WT N= 6 mice. V_321_L N=5 mice, p= 0.004 (**), Mann-Whitney Rank Sum Test U=0). **(D) Left,** Image from a WT DG showing neural stem cells infected with a replication-deficient GFP-expressing retrovirus via stereotaxic injection, followed by 14 days of incubation to label NPs and their progeny. The identities of GFP+ cells were determined by a combination of morphology, Dcx immunoreactivity, and Nissl staining. Neural progenitors (NPs) were identified by the lack of radially oriented processes with or without Dcx immunoreactivity (open arrowheads). Immature granule cells (iGCs) were identified by their radially oriented neurites, Dcx+ and Nissl+ staining (filled, yellow arrowheads). Glial cells were identified by their star-like morphology and lack of Dcx immunoreactivity and Nissl staining (yellow arrows). **Right,** Fate specification was quantified as proportion of GFP+ cells that remained radial glial-like stem cells, NPs, or committed to neuronal or glial fates. 5-14 50-μm sections were quantified from each animal, depending on infection efficiency (WT N=3 mice, V_321_L N=4 mice; Two-way ANOVA Genotype x Cell Fate p=0.0003 F (3, 20) =9.864. NP, WT vs. V_321_L Bonferroni-corrected p=0.003 (**). iGC, WT vs. V_321_L Bonferroni-corrected p=0.007 (**). RGL & Glial comparisons between genotypes were not significant p>0.9999).

The discrepancy between decreased proliferation and unchanged neuronal production in the V_321_L DG could be explained by disruptions to the balance between self-renewal maintenance and postmitotic neuronal differentiation, both of which are processes predicted to be disrupted by the V_321_L mutation based on the transcriptomic data (**Table 5-2)**. Therefore, we asked whether cell cycle dynamics and/or fate commitment by neural stem cells were abnormal in the V_321_L DG.

To measure the rate of cell cycle reentry, cells in S phase of the cell cycle were first labelled by systemic EdU injection. 24 hours after EdU injection, we collected brain sections and stained them for Ki67 expression. We calculated the overall rate of cell cycle reentry as the proportion of total EdU+ cells that were also Ki67+. We observed a statistically significant decrease in cell cycle re-entry in the V_321_L DG compared to WT DG (**Figure 7C**; p= 0.0043, Mann-Whitney Rank Sum Test). Therefore, the self-renewing pool of neural progenitors was depleted faster, generating more postmitotic cells in the V_321_L mutant dentate gyrus.

Given the higher rate of cell cycle exit in V_321_L DG, we asked if fate programming was affected. V_321_L and WT DGs were injected with a replication-deficient retrovirus encoding GFP, which only stably infects cell during mitosis. After 14 days, brains were collected for identification of GFP+ progeny. The identity of GFP+ cells were determined by a combination of morphology, Dcx immunoreactivity, and Nissl staining. Neural progenitors were identified by short, tangential processes and the lack of radially oriented processes, with or without Dcx immunoreactivity (**Figure 7D**, open arrowheads). Immature GCs (iGC) were identified by their radially oriented neurites and Dcx+ Nissl+ staining (**Figure 7D**, closed arrowheads). Glia (Astrocytes/Oligodendrocytes) were identified by their star-like morphology, extensive branch ramifications, and lack of Dcx immunoreactivity and Nissl staining (**Figure 7D**, yellow arrows).

The results revealed a decrease in the proportion of neural progenitors in the V_321_L DG compared to WT DG (**Figure 7D**; Two-way ANOVA Genotype x Cell Fate p=0.0003 F (3, 20) =9.864. NP, WT v. V_321_L Bonferroni-corrected p=0.0026), consistent with the observed depletion of the proliferative pool (**Figure 7A**) and increased rate of cell cycle exit (**Figure 7C**). Interestingly, fate commitment in the V_321_L DG was disproportionately biased towards the neuronal fate program compared to glial fate (**Figure 7D**; iGC, WT v. V_321_L Bonferroni-corrected p=0.0069). In fact, the proportion of glia generated by the V_321_L neural stem cells was not significantly different between V_321_L and WT DG (Glia, WT v. V_321_L p>0.9999).

Thus, we found depletion of the proliferative neural stem cell pool, increased cell cycle exit, and disproportionate production of neurons in the V_321_L DG in agreement with predictions from our transcriptomic analyses (**Figure 5**). These findings point to an important role for Nrg1 nuclear back signaling for neural stem cell maintenance and fate commitment.

### The Nrg1 V_321_L mutation compromises maturation of granule cells in the dentate gyrus

Nrg1 nuclear back-signaling is important for dendrite arborization and spine production (Chen, Hancock et al. 2010, Fazzari, Snellinx et al. 2014). V_321_L cortical neurons showed loss of ErbB4-induced dendrite growth (**Figure 2E**), inhibition of gamma secretase resulted in a dramatic reduction in dendrite growth (**Figure 3C**), and nuclear back signaling regulated genes were predicted to enhance the development of neurites (**Figure 6C**). Our RNASeq analysis of the V_321_L DG also showed dysregulation of genes involved in dendrite development (**Table 5-2**). Therefore, we first examined the dendrite morphology of newborn dentate GCs.

Qualitative observations from Dcx immunohistochemistry revealed decreased dendritic complexity of adult-born immature neurons in V_321_L DG (**Figure 8A**). To systematically quantify mature granule cell dendritic arbors, we used Golgi impregnation to visualize neuronal morphology and only analyzed dendrites of neurons with typical granule cell morphologies (see methods for details on criteria). **Figure 8B** shows representative tracings of granule cells from each genotype. Sholl analysis revealed significant increases in V_321_L dendritic length proximal to the soma (**Figure 8C**; at 80μm away from soma Bonferroni-corrected p=0.052, Two-way RM ANOVA with a significant (Genotype x Distance from soma) interaction p=0.035).

**Figure 8:**
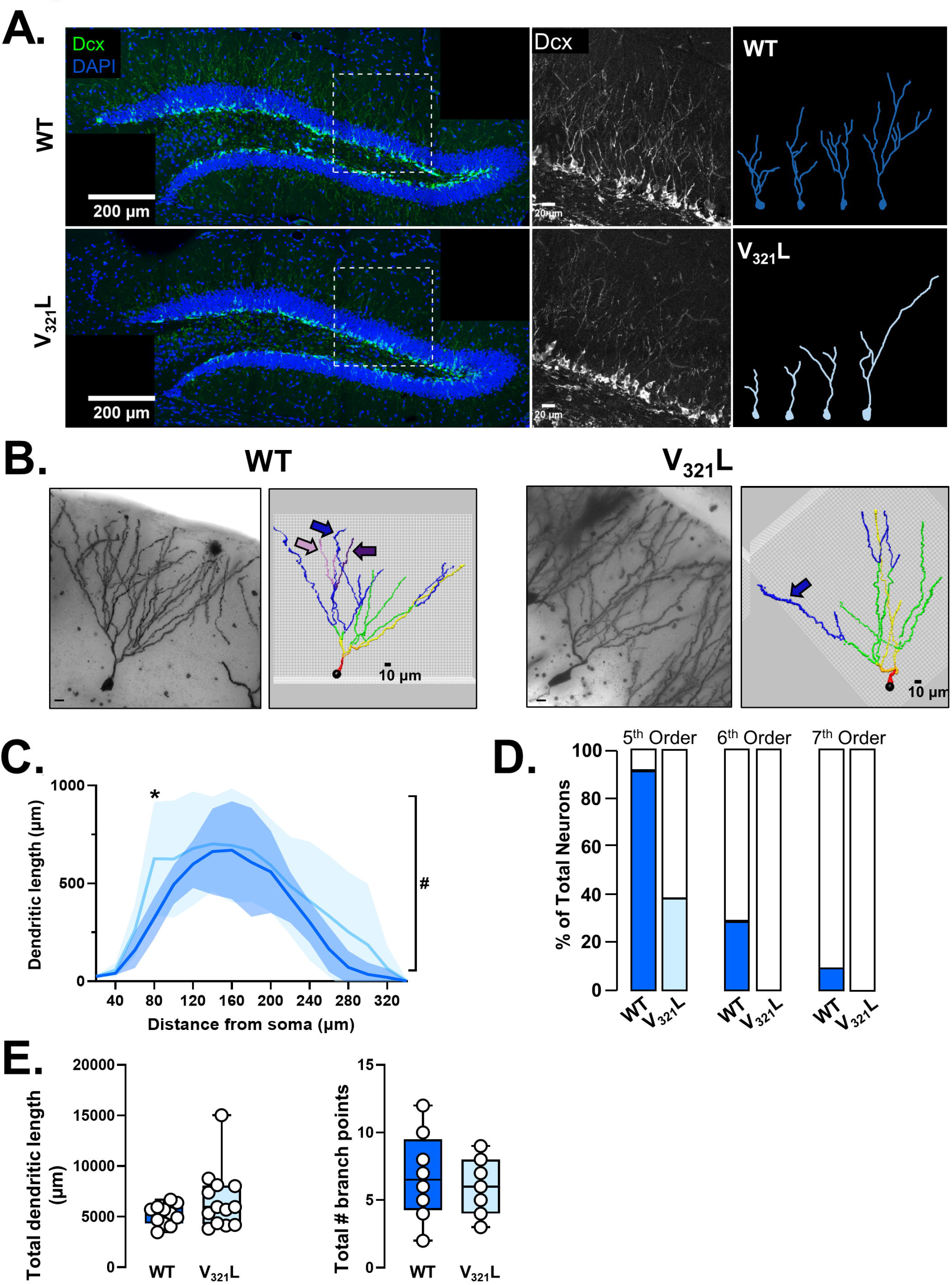
Granule cells produced in the V_321_L DG display aberrant dendritic arborization. **(A) Left**, Doublecortin (Dcx - green) immunohistochemistry in the P72 DG along with DAPI counterstain (blue), with high magnification of the dotted white boxes in the middle. **Right,** Representative tracings of Dcx+ cells in WT and V_321_L DG. **(B)** Representative images and tracings of Golgi-impregnated DG granule cells (GCs) from ∼P60 WT and V_321_L. Neurites are color coded by branch order (1°= red, 2°= orange, 3°= yellow, 4°= green, 5°= blue, 6°= purple, 7°= pink). A representative neuron is shown for each genotype. Arrows following the color code outlined above highlight the absence of higher order branching in V_321_L GCs. **(C)** Length of dendritic segments within each zone were summed. Sholl analysis reveals difference in distribution of dendritic lengths as a function of distance from the soma in V_321_L mice. Shaded region around the line represents standard deviation. Data is compiled from 11 neurons from 5 WT (dark blue) animals and 13 neurons from 5 V_321_L (light blue) at ∼P60. There was a significant difference in the sholl analysis profile between WT and V_321_L neurons (Mixed effects model Two-way RM ANOVA Genotype x Distance from soma p=0.03(#), F (16, 352) =1.763; WT vs. V_321_L at 80μm away from soma FDR corrected q=0.05 (*)). **(D)** Proportion of neurons in which 5^th^, 6^th^, and 7^th^ order branching are present. V_321_L granule cells had fewer branches at the higher orders of branching (5^th^ order, p=0.01; 6^th^ order, p= 0.08; 7^th^ order, p=0.46; Fisher’s exact test). (WT (dark blue) N=11 neurons/ 5 mice, V_321_L (light blue) N=13 neurons/ 5 mice). **(E)** (**Left**) Total dendritic length of Golgi-impregnated granule cells quantified using NeuronStudio. Total dendritic length trends greater in V_321_L mice, though not statistically significant (p= 0.25, Wilcoxon rank sum test) (**Right**) Total number of branch points trends lower in V_321_L DGs compared to WT counterparts, although the difference is not statistically significant (p= 0.45, Welch’s t-test) (wildtype N=11 neurons/ 5 mice, V_321_L N=13 neurons/ 5 mice).

Dendrite complexity appeared to be compromised in GCs sampled from V_321_L DGs. In WT neurons most of the dendritic length was distributed in 4^th^ and 5^th^ order, whereas in V_321_L GCs, the peak of dendritic length was found in 4^th^ order branches with less length distributed in 5^th^ order branches (data not shown). In fact, there was a statistically significant decrease in the proportion of GCs that possessed 5^th^ order branching in V_321_L GCs, and while a small percentage of WT GCs possessed 6^th^ and 7^th^ order branches, these high order branches were not observed in V_321_L GCs (**Figure 8D**, 5th order: p= 0.0131, 6th order: p= 0.0815, 7th order: p= 0.458, Fisher’s exact test). Total dendritic length and number of branch points were not significantly altered in the V_321_L DG (**Figure 8E; total length**: p=0.2518 Mann-Whitney Rank Sum Test, U=51; **branch points**: p=0.6035 Welch’s t-test t=0.5331, df=12.23).

Altogether, examination of dentate GCs revealed alterations in the dendrites in both immature and mature V_321_L GCs, especially in distal dendrites. Thus, diminished Nrg1 nuclear back-signaling results in persistent and cumulative effects on dendrite growth and complexity across development.

### Cell autonomous rescue of dendritic growth by the Nrg1 ICD

Nrg1 nuclear back signaling requires gamma secretase activity (**Figure 2C**), and gamma secretase activity is necessary for proper dendrite development in GC-enriched hippocampal cultures (**Figure 3C**). Finally, dysregulated gene expression in the V_321_L DG are enriched for genes involved in dendrite morphogenesis and mutant mice show altered GC dendritic morphology (**Table 5-2** and **Figure 8**). Thus, we hypothesized that the gamma secretase dependent signal required for dendritic growth might be the nuclear Nrg1 ICD. To test this hypothesis, we cultured P4 hippocampal neurons in the presence of DAPT from DIV1-14 as described earlier. To one group of DAPT treated neurons and one group of vehicle treated neurons, we added AAV_9_ particles to deliver an expression construct for the Nrg1 ICD-Flag (AAV-ICD) (**Figure 9A, left**). Neurons were cultured for 14 days after which dendritic growth was assessed via MAP2 and successful re-expression of the nuclear ICD was confirmed via Flag staining and evaluating nuclear localization of the Flag-tagged ICD (**Figure 9A, right**). Consistent with the results shown in Figure 3, we found that DAPT treatment significantly disrupted dendritic growth, but this effect was abolished by re-expression of the nuclear Nrg1 ICD. Importantly, in cultures treated with the AAV-ICD, neurons that lacked detectable levels of the Nrg1 ICD in the nucleus were not rescued from the effects of DAPT (**Figure 9A right, and Figure 9C**; Kruskal-Wallis test p=0.0001 KW= 22.94; DAPT vs. DAPT+AAV-ICD p=0.03; DAPT vs. Veh.+AAV-ICD p= 0.004; DAPT+AAV-ICD vs. DAPT+AAV-ICD (non-expressing) p=0.0045; DAPT+AAV-ICD (non-expressing) vs. Veh+AAV-ICD p= 0.0007; DAPT vs. DAPT+AAV-ICD (non-expressing) p>0.9999).

**Figure 9:**
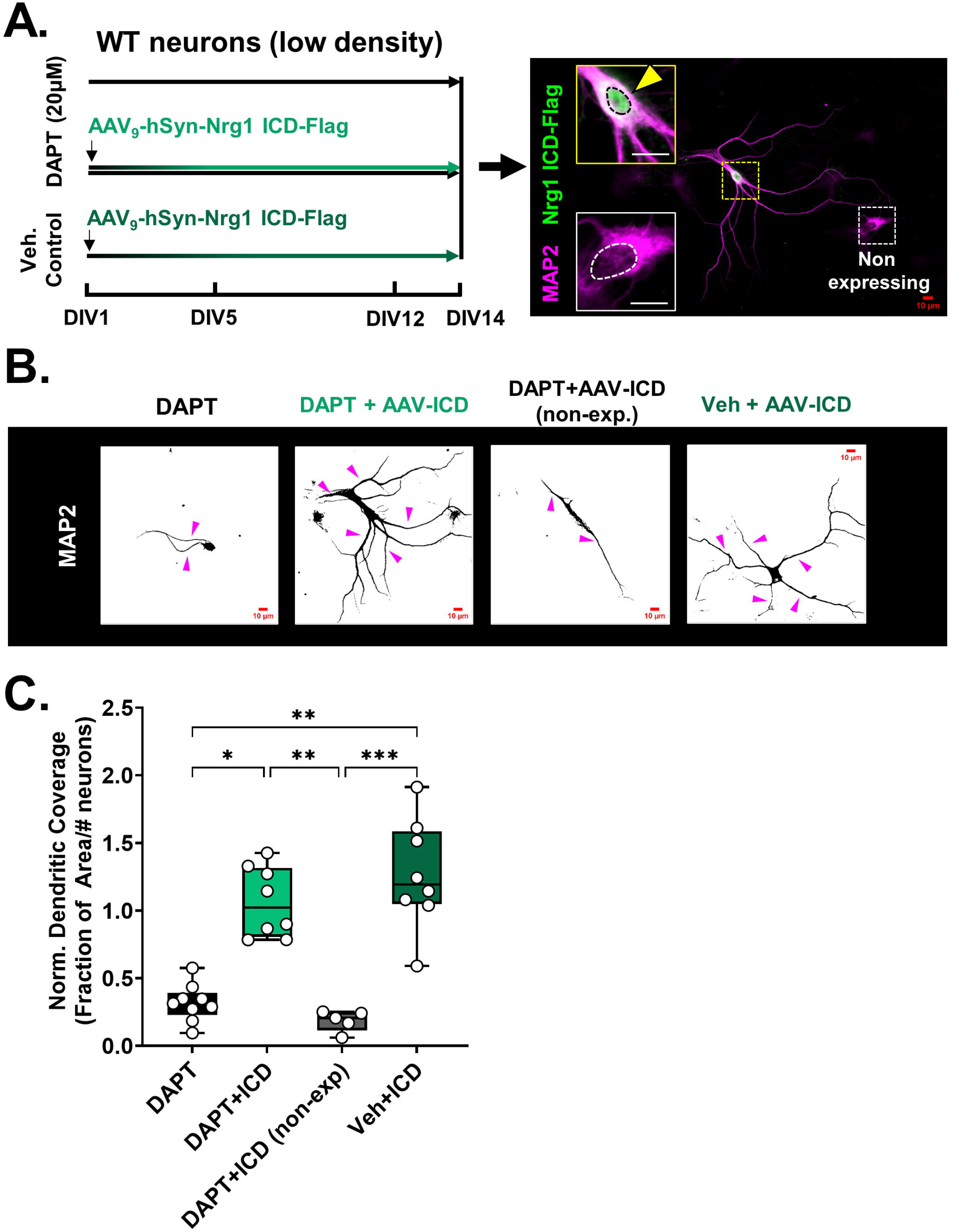
Gamma secretase-dependent dendrite growth requires the nuclear Nrg1 ICD. **(A)** (**Left**) Schematic of the various treatment regimens. Vehicle controls received equal volumes of DMSO diluted in culture media. DAPT treatment was started from DIV1 and continued until DIV14. One of the DAPT treatment groups and the vehicle control group, on DIV1, were also incubated with AAV9 particles to deliver an expression construct for the Nrg1 ICD (flag-tagged) under the control of the human synapsin (hSyn) promoter. All platings were collected at DIV14, fixed and stained for MAP2 and Flag. (**Right**) Representative image of a culture treated with DAPT from DIV1-14 and infected with AAV_9_-hSyn-Nrg1ICD-Flag, stained for MAP2 and Flag on DIV14. The image shows two neurons in the same plating, one that is expressing the ICD and another that is non-expressing. Insets show each of these cells at higher magnification and the dotted line delineates the nucleus. The yellow arrowhead indicates Nrg1 ICD expression. Scale bar= 10µm.**(B)** Representative images of MAP2 staining at DIV14 for each of the treatment conditions described in **panel A**. Magenta arrowheads indicate MAP2+ neurites. Scale bar= 10µm. **(C)** Disrupted dendrite outgrowth by gamma secretase inhibition was rescued by expression of the nuclear Nrg1 ICD. Quantification of fraction of area covered by MAP2 staining normalized to the number of neurons in the imaging field. Inhibition of gamma secretase significantly reduced dendritic growth, which was rescued by re-expression of the nuclear Nrg1 ICD. Kruskal-Wallis test p=<0.0001 KW=22.94. DAPT vs. DAPT+ICD p=0.03 (*), DAPT vs. Veh+ICD p= 0.004 (**), DAPT+ICD vs. DAPT+ICD (no exp) p=0.0045 (**), DAPT+ICD (no exp) vs. Veh+ICD p=0.0007 (***). N=8-10 platings from 6-8 mice from 2 litters per genotype for each condition.

The dendritic defects originating from gamma secretase hypofunction can be rescued in a cell autonomous manner by re-expression of the nuclear Nrg1 ICD, however, we posit that the endogenous source of this nuclear ICD is a consequence of axon-dendrite interactions during synaptogenesis (**Figure 4**).

### Nrg1 V_321_L mutant mice show sensorimotor gating deficits and a hyperresponsive auditory startle reflex

Orthologs of schizophrenia-associated genes were found to be dysregulated in the V_321_L DG (**Table 5-3**). The V_321_L mutation in Nrg1 was discovered using a family-based association test where it was found to be over-transmitted to offspring with psychosis and schizophrenia (Walss-Bass, Liu et al. 2006). Additionally, both Type III Nrg1 heterozygous mice and mice deficient in gamma secretase show sensorimotor gating deficits, an endophenotype of psychotic disorders (Chen, Johnson et al. 2008, Dejaegere, Serneels et al. 2008). Thus, we asked whether the V_321_L mice showed sensorimotor gating deficits by assessing pre-pulse inhibition (PPI) of the auditory startle reflex. We first subjected WT and V_321_L mice to 60 trials of a 115dB startle stimulus to measure the startle amplitude and assess habituation of the startle response (**Figure 10A**). We found that neither WT nor V_321_L mice showed significant habituation of the startle reflex (**Figure 10A**; Two-way RM ANOVA, Effect of Trial number p=0.3). However, V_321_L mice consistently showed significantly larger startle responses compared to WT mice (**Figure 10A**; Two-way RM ANOVA, Effect of Genotype p=0.005). Next, the mice were subjected to 160 trials consisting of startle stimulus delivered alone or preceded by a pre-pulse of varying amplitudes and varying delay (**Figure 10B**). WT mice showed significantly higher PPI on trials consisting of either 75dB or 85dB pre-pulses compared to either 68dB or 70dB trials, neither of which elicited PPI (**Figure 10B**; Two-way RM ANOVA, Bonferroni corrected p<0.05). V_321_L mice only showed statistically significant increase in PPI on 85dB trials compared to other trials, however, PPI was significantly impaired compared to WT mice (**Figure 10B**; Two-way RM ANOVA, Genotype x Trial Type p<0.0001). Additionally, V_321_L mice showed significantly higher startle responses on the non-prepulse trials interspersed throughout the PPI trial block (WT v. V_321_L p=0.03 Welch’s t-test t=2.362, df=15.91). Upon comparing startle responses during non-prepulse trials to those with pre-pulses, we noticed a general trend towards sensitization of the startle response in the V_321_L mice compared to WT mice (**Figure 10C, left**). We found that a larger proportion of V_321_L mice show sensitization on a greater number of trials irrespective of pre-pulse intensity, whereas WT mice only ever showed sensitization during the mild pre-pulse trials (68-70dB) and always showed startle suppression on the relatively stronger pre-pulse trials (75 and 85dB) (**Figure 10C, middle and Figure 10C, right** WT v. V_321_L p=0.05 Welch’s t-test t=2.147, df=14.39).

**Figure 10:**
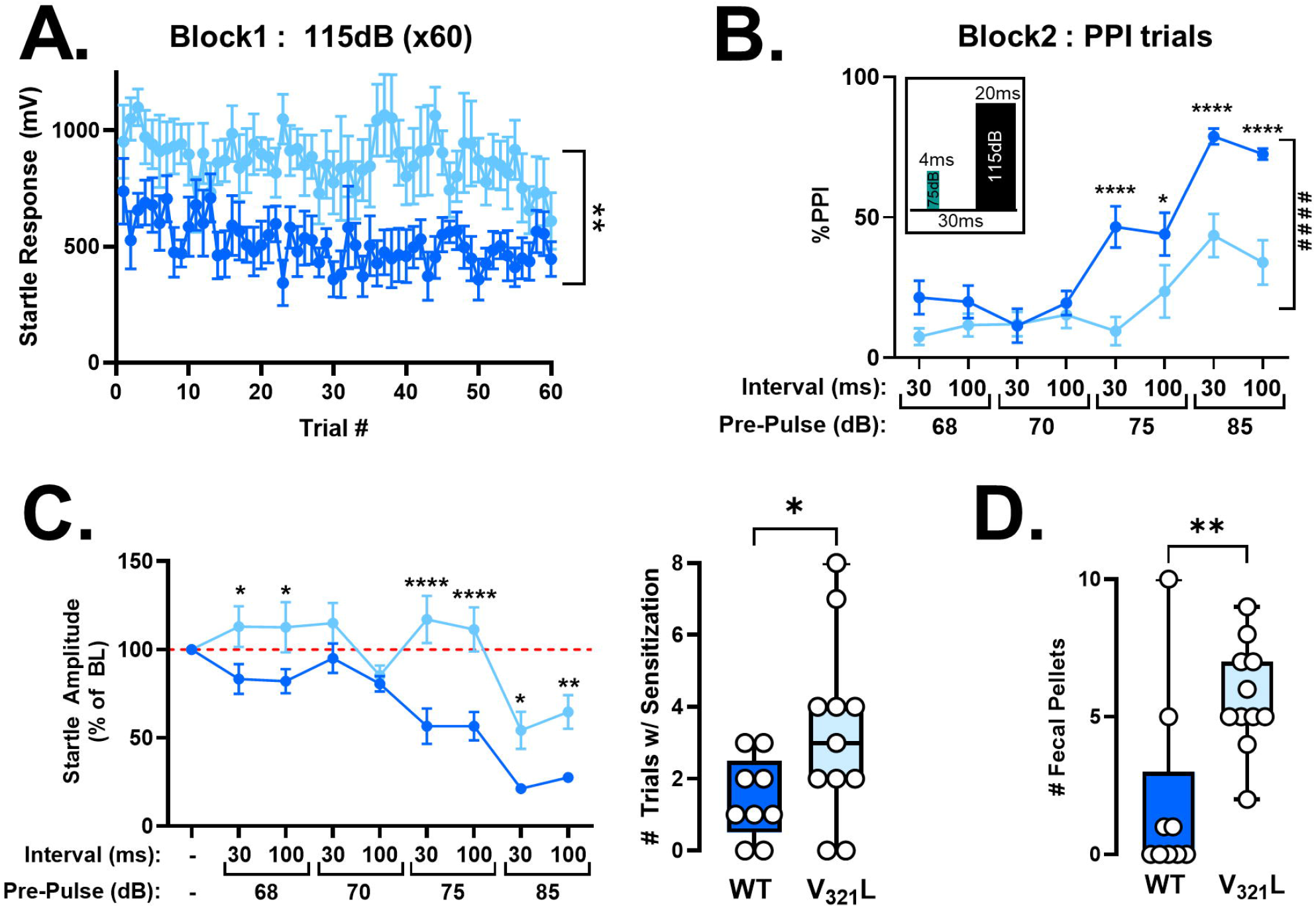
V_321_L mutant mice show sensorimotor gating deficits. **(A)** Startle responses during Block 1 trials, comprised of 60 startle stimuli (115dB). Neither WT (dark blue) nor V_321_L (light blue) mice showed significant habituation of the startle response during the testing session. However, V_321_L mice showed a significantly larger startle response (**); Two-way RM ANOVA Trial x Genotype p=0.40, Trial p=0.27, Genotype p=0.005 F (1,18) = 10.45. Data represented as Mean±SEM. N=9 mice (WT) and 11 mice (V_321_L). **(B)** V_321_L mice showed significantly lower PPI compared to WT mice; Two-way RM ANOVA Trial Type x Genotype p<0.0001 (####) F (7,126) = 5.387, Trial Type p<0.0001, Genotype p=0.0026. Multiple comparisons (WT vs. V_321_L; Bonferroni corrected): 75dB-30ms-115dB p<0.0001 (****), 75dB-100ms-115dB p=0.02 (*), 85dB-30ms-115dB p<0.0001 (****), 85dB-100ms-115dB p<0.0001 (****). Data represented as Mean±SEM. N=9 mice (WT) and 11mice (V_321_L). Inset shows a schematic of a representative PPI trial consisting of a 4ms 75dB pre-pulse and a 20ms 115dB startle stimulus separated by 30ms. The pre-pulse intensities and intervals between pre-pulse and startle pulse was varied; different combinations are displayed on the x-axis. **(C)** V_321_L mice showed sensitization of the startle response. (**Left**) Plot shows startle amplitude normalized to baseline startle response (BL), which is the average of the startle response during startle alone trials in Block 2. Red dashed line delineates 100%, representing no suppression of startle response. An increase beyond baseline startle (red dashed line) indicates a sensitized response; Two-way RM ANOVA Trial x Genotype p<0.0001 F (8,144) = 4.977, Trial Type p<0.0001, Genotype p=0.009. Data represented as Mean±SEM. 9/11 V_321_L and 4/9 WT mice showed sensitization on at least one of the trials. (**Right**) V_321_L mice showed sensitized startle responses on more trials than WT mice; Welch’s t-test (two-tailed) p=0.05 t=2.147, df=14.39. N=9 mice (WT) and 11mice (V_321_L). **(D)** V_321_L mice showed increased defecation compared to WT mice consistent with a more anxious behavioral state; Mann-Whitney test p=0.006, U=15. N=9 mice (WT) and 11mice (V_321_L).

We also measured number of fecal pellets as a measure of anxiety (Hall 1934). Mice were handled and acclimated to the testing environment for three days and then subjected to the PPI testing protocol on two consecutive days. WT mice showed little to no defecation during the final PPI testing, however, V_321_L mice continued to defecate at significantly higher rates than WT mice indicating a potentially higher anxiety-like behavioral state (**Figure 10D**; WT v. V_321_L p=0.006 Mann-Whitney Rank Sum Test, U=15).

### Dysregulation of a functionally connected schizophrenia susceptibility gene network in the V_321_L DG

Transcriptomic enrichment analyses combined with empirical studies shown thus far strongly implicated the Nrg1 ICD in regulating a functionally connected network of genes involved in neuronal development. The enrichment of orthologs of schizophrenia-associated genes within the dysregulated genes in the V_321_L DG raised the possibility that the schizophrenia-associated genes might also function within this larger network of genes regulating neuronal development. To gain further insight into how schizophrenia-associated DEGs in the V_321_L DG, might regulate neuronal development, we first asked if the schizophrenia-associated DEGs in the V_321_L DG functionally interact with the larger set of all the other DEGs in the V_321_L DG. To do this, we created a merged list of dysregulated genes in the V_321_L DG whose human orthologs are associated with psychosis, schizophrenia, sporadic schizophrenia, or schizoaffective disorder, under the umbrella term of “schizophrenia spectrum disorders” (SCZ) yielding a total of 67 genes (**Figure 11A**; enrichment p value =5e-7; **Table 11-1**). We then grew a network (see methods) by providing the rest of the significantly DEGs in the V_321_L DG seeking to find genes in our dataset that have not been explicitly identified as schizophrenia-associated but might have direct functional interactions with dysregulated schizophrenia-annotated genes. This resulted in a network with 322 nodes and 371 edges, which we called the “SCZ+” network. 49 of the 67 SCZ genes were incorporated into this SCZ+ network. (**Figure 11B, right. Table 11-1**). Thus, the bulk (73%) of the schizophrenia-associated DEGs do interact within a larger network of DEGs, which as previously shown, are involved in various neurodevelopmental processes (**Figure 5 and Table 5-2**). Next, we analyzed the topological properties of this network to derive further insight into the nature of these interactions aiming to further distill the role SCZ genes might play within this network.

**Figure 11:**
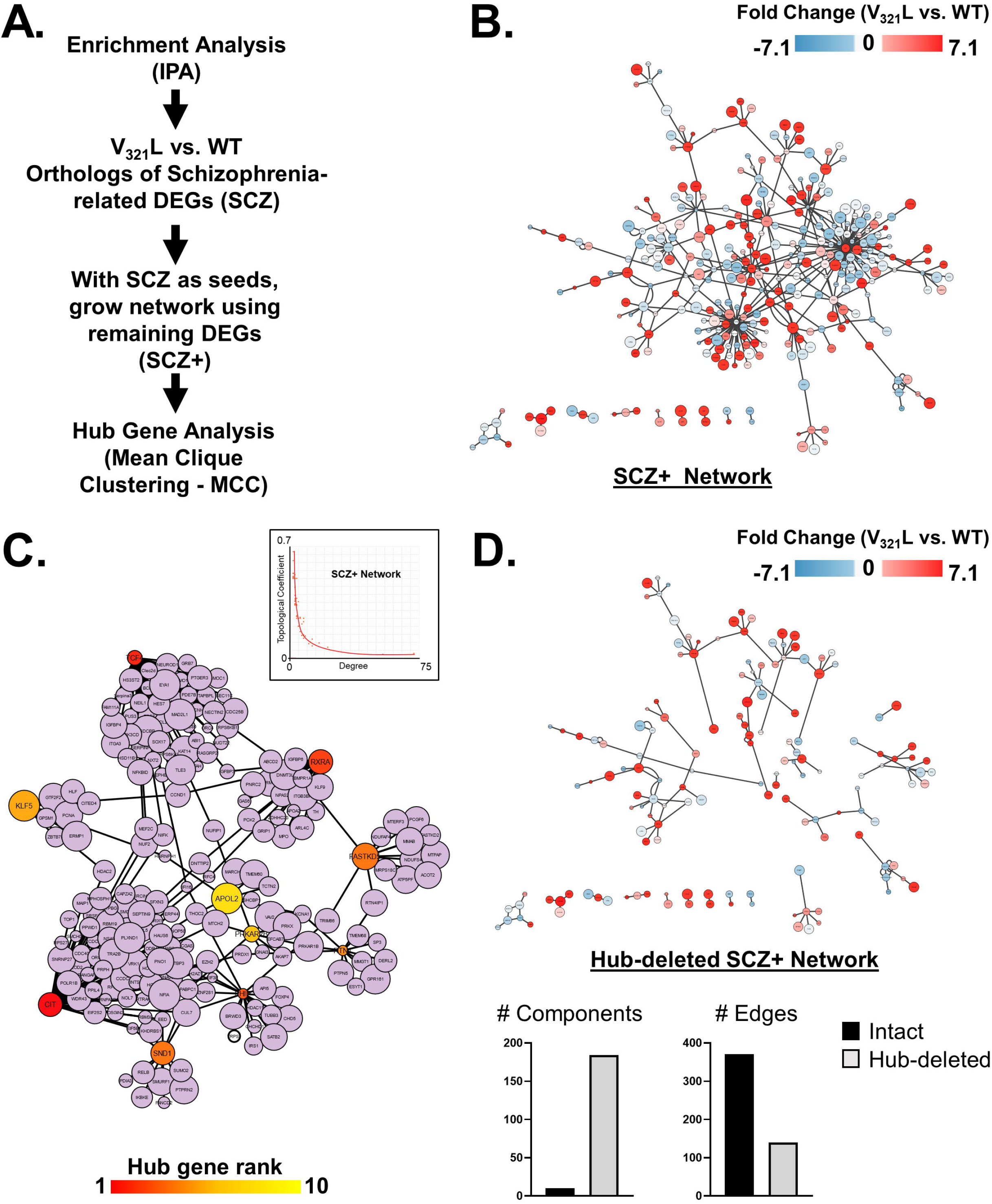
Dysregulation of a schizophrenia-susceptibility gene network in the V_321_L DG. **(A)** Workflow outlining the generation and analysis of a schizophrenia-susceptibility gene network. **(B)** Orthologs of genes associated with schizophrenia spectrum disorders (Figure 5D-IPA analysis) were found to be dysregulated in the V_321_L DG (**Table 5-3**). A network was grown by seeding schizophrenia-related dysregulated genes (V_321_L vs. WT) along with the remaining set of differentially expressed genes between V_321_L and WT DG, in IPA. We labeled this network – <u>SCZ+</u>. Node colors represent fold change in expression (-7.1 blue to 7.1 red) according to heatmap. This network was subjected to hub gene analysis using the mean clique clustering (MCC) method. Also see **Extended Data Table 11-1, 11-2**. **(C)** Network display of 10 hub genes identified by MCC analysis of the SCZ+ network, along with their first-degree neighbors (lavender colored nodes) shows a hub-spoke connectivity with schizophrenia-related genes forming hubs. Hub node color reflects MCC rank. Inset shows a plot showing the relationship between the degree and topological coefficient of every node in the SCZ+ network. The inverse relationship indicates the presence of potential hubs in the network. **(D) Top,** Hub genes identified in panel C were deleted from the SCZ+ network, disconnected nodes were removed from the network and the resulting network is displayed. **Bottom,** Quantification of number of connected components and edges in the SCZ+ network and Hub-deleted SCZ+ network. Node colors represent fold change in expression (-7.1 blue to 7.1 red) according to heatmap.

Analysis of the expanded schizophrenia-susceptibility network (SCZ+) implicated a “hub and spoke” topology. Hub nodes interacted preferentially with a few other nodes as opposed to with each other (i.e. the degree of a node and its topological coefficient were inversely related) (**Figure 11C inset**). We identified 10 hub nodes using mean clique clustering (MCC), 9 of which were part of the primary schizophrenia-susceptibility network (**Figure 11C**). The 10 hubs shared no edges with each other thereby creating 10 modules (red (higher rank) to yellow (lower rank) MCC Top10 nodes). We visualized the first-degree neighbors of these hub genes and found that each of the hub genes coordinated its own mini network as predicted from the network topology (**Figure 11C**). To validate the status of the hub genes as hubs within the SCZ+ network, we reconstructed the SCZ+ network shown in **Figure 11A** after removing the hub genes from the network. Exclusion of these 10 hub genes resulted in fragmentation of the network indicated by the drastic increase in the number of components and decrease in the number of edges (**Figure 11D**). Thus, the SCZ genes not only interact with a larger network of dysregulated genes, but a subset of them (9 out of the original 67) are essential for organizing the network.

Next we performed Gene Ontology analyses on the 49 SCZ genes and the 271 DEGs that became part of SCZ+ network. The SCZ genes were enriched for functions related to synaptic transmission (adj.p=4.02e-8) and were enriched for genes whose products are localized to synaptic membranes (SynGO-adj.p<0.01). Intriguingly, none of the hub genes (90% of which were part of the original schizophrenia-associated gene list) were associated with synaptic transmission. The 271 DEGs that comprised the SCZ+ network were also enriched for the annotation-transcriptional regulation (adj.p=0.02) (driven by 52 of the 271 genes) and were enriched for genes whose products are localized to the nucleus (adj.p=0.00002) (**Table 11-2**). These data show that while schizophrenia associated DEGs are enriched for genes involved in synaptic function, hubs that embed these core genes into cellular signaling networks are not obvious mediators of synaptic function.

### Nrg1 nuclear back-signaling regulates functionally related gene modules

Transcriptional regulator (TR) analysis showed that DEGs in the V_321_L DG can be regulated by a small set of transcriptional regulators (e.g. REST, PRC2, E2F4, etc.). TR analysis also showed that DEGs regulated by the same regulators were functionally related (**Tables 5-4 and 5-5**). This result raises the possibility that the Nrg1 ICD transcriptionally regulates functionally related genes. To investigate this, we performed weighted gene correlation network analysis (WGCNA) to identify gene co-expression modules associated with the V_321_L mutation (**Figure 12A and 12B**). WGCNA identified 2 modules in which expression of genes was statistically significantly correlated (**Brown module: mouseUP module**; p=0.0002, corr.coefficient (LL genotype)= 0.9) or anticorrelated (**Turquoise module: mouseDOWN module**; p=0.009, corr.coefficient (LL genotype)= -0.74) with the mutant genotype (**Figure 12C**). We performed permutation analyses to validate the correlation coefficients and p-values and found that randomized module membership could not achieve the same values as observed (**Figure 12D**).

**Figure 12:**
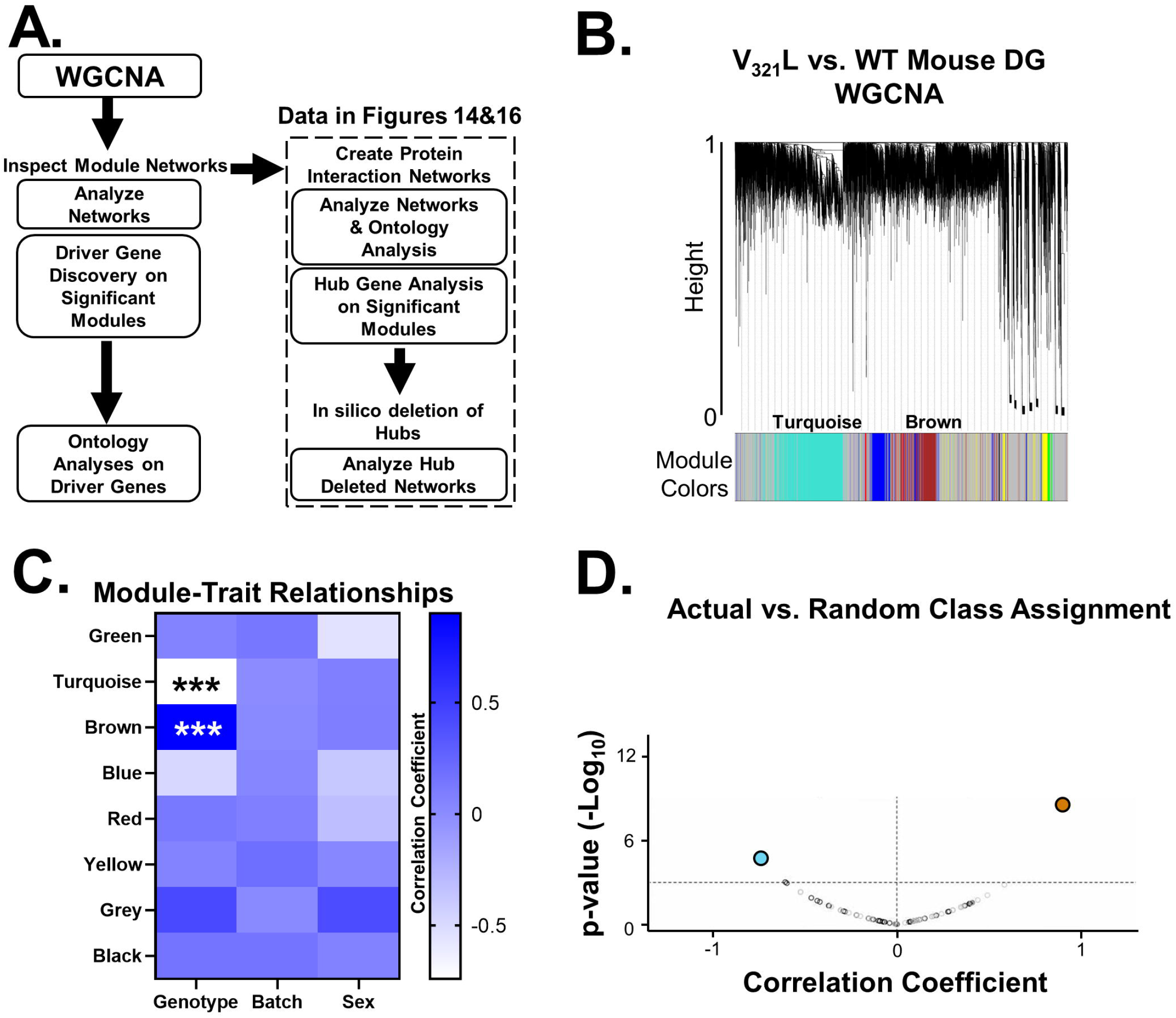
Weighted Gene Correlation Network Analysis (WGCNA) of V_321_L mouse DG. **(A)** Flowchart outlining the analysis strategy following WGCNA module discovery. Protein interaction (PI) networks constructed from WGCNA modules are shown in Figure 14. **(B)** Module detection (WGCNA) via dynamic tree cut from the WT vs. V_321_L DG transcriptome. The detected modules are represented by colors below the dendrogram. The two statistically significant modules (Turquoise and Brown) are labeled. **(C)** Heatmap showing module-trait relationships between detected modules in panel A and the V_321_L genotype, sequencing batch (technical factor), and sex of the animal. The color indicates the correlation coefficient according to the legend. Only two modules were statistically significantly associated with genotype and not with sequencing batch. We dubbed the Turquoise module as the **mouseDown module** (p=0.009) and the Brown module as the **mouseUp module** (p=0.0002) for a more intuitive nomenclature. None of the modules were associated with chromosomal sex of the mice. **(D)** Permutation analysis of module discovery-gene memberships were permuted 40 times. None of the permuted modules had equivalent correlation coefficients and statistical significance as the observed modules. The grey open circles represent permuted turquoise modules and black open circles represent permuted Brown modules.

Next, we asked whether the genes whose expression was significantly associated with the mutant genotype were also important members of these modules as this would warrant further investigation into the gene networks that comprise these modules. To do this, we examined the relationship between module membership (MM – co-expression between each gene in a module with the module eigengene) and gene significance to the mutant genotype (GS – correlation of gene expression with genotype) (**Figure 13A, insets**). Both modules showed significant correlation between these two metrics indicating that the genes associated with (dysregulated by) the mutant genotype were proportionately important to the networks within each module (**Fig 13A Top (inset),** mouseUP: corr=0.79, p<0.0001 & **Figure 13A Bottom (inset),** mouseDOWN: corr=0.53, p<0.0001). We identified genes with the 95^th^ percentile of module membership scores within each module as putative driver genes for the network. These genes were then removed from the network. Since the network was weighted, we predicted that loss of driver genes would bias the network to lower weighted edges thereby destabilizing the network. Indeed, loss of the identified driver genes created a significant left shift in the weight distributions in both modules (**Figure 13A, Top.** mouseUP: KS test p=2.2e-16 and **Bottom.** mouseDOWN: KS test p=2.2e-16).

**Figure 13:**
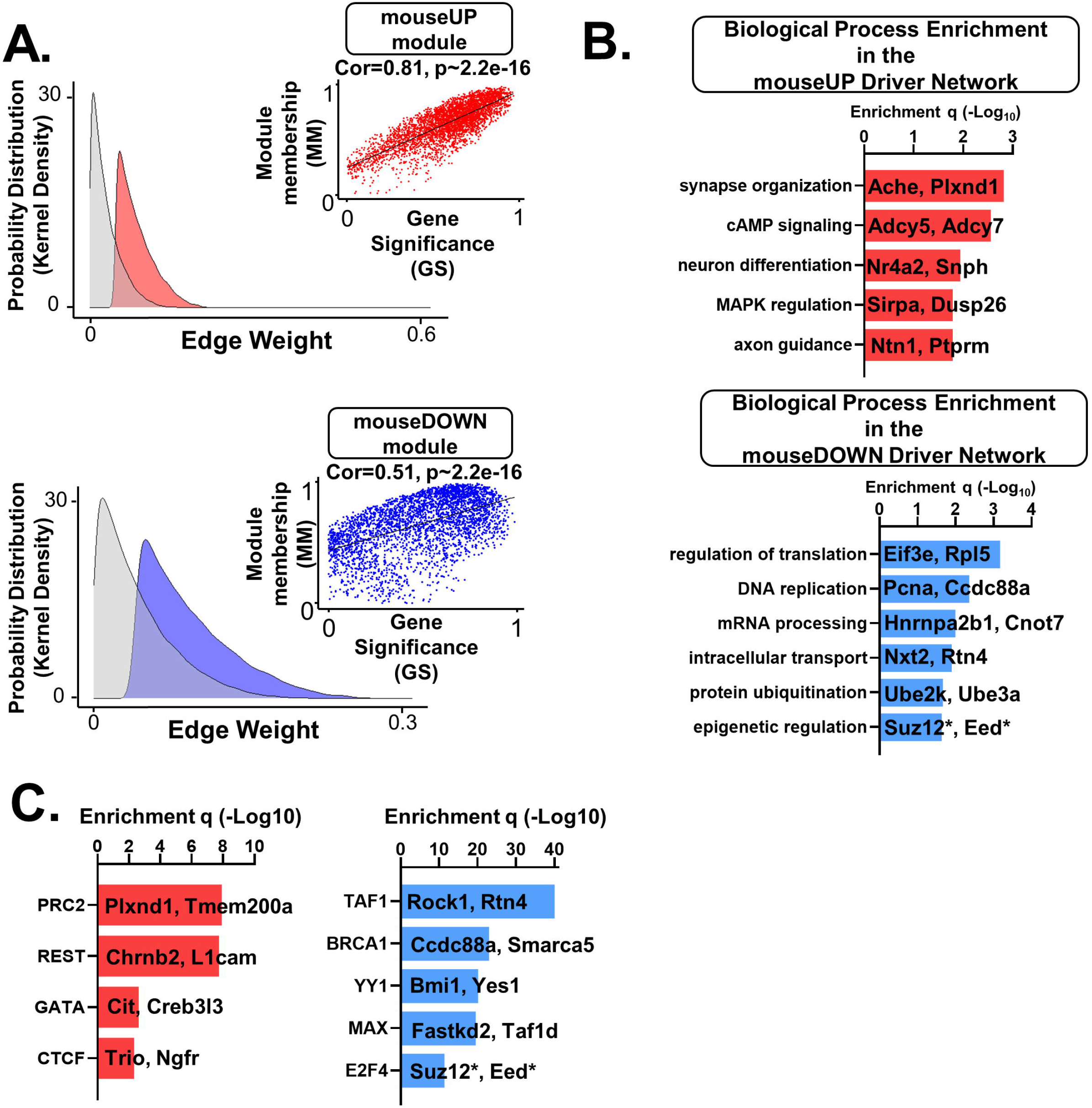
Analysis of driver genes and driver gene networks from the V_321_L mouse WGCNA. **(A)** Deletion of selected driver genes from the WGC network alters the weight distribution of the network indicating their significance within the network. **(Top)** MouseUP (intact (red) vs. driver deleted (grey): Kolmogorov Smirnov D=0.05 p∼2.2e-16) and **(Bottom)** MouseDOWN (intact (blue) vs. driver deleted (grey): Kolmogorov Smirnov D=0.59 p∼2.2e-16). **Insets** show correlation between gene significance (GS) and module membership (MM) for both mouseUP and mouseDOWN modules. Correlation analysis reveals a strong relationship between GS & MM for the mouseUP module (Pearson corr. 0.81; p=2.2e-16) indicating their significance to the dysregulated V_321_L DG transcriptome and importance within the mouseUP module. Correlation for the mouseDOWN module was not as strong but still revealed a significant relationship between GS & MM (Corr. 0.5; p=4.2e-16); Driver genes were selected as ones having the highest MM (Top 95^th^ percentile). **(B)** Ontology analysis showing biological processes enriched within the driver gene network. **(Top, red)** Biological Processes implicated in the mouseUP driver gene network. **(Bottom, blue)** Biological processes implicated in the mouseDOWN driver gene network. Asterisks highlights the presence of PRC2 core components in the mouseDOWN module. **(C)** Predicted transcriptional regulators for the driver genes from the ChEA and ENCODE databases reveal polycomb repressor complex2 (PRC2) as a strong candidate in regulating genes in the mouseUP module and TAF1 for the mouseDOWN module. Asterisks highlights the presence of PRC2 core components in the mouseDOWN module predicted to be regulated by E2F4.

To understand what functions, if any, were overrepresented by the driver genes, we performed Gene Ontology analyses on the driver genes and found that the mouseUP driver genes were enriched for predominantly axonal biased functions – synapse organization, cAMP signaling, axon guidance, along with others such as neuronal differentiation and negative regulation of MAPK signaling (**Figure 13B, Top)**. The mouseDOWN driver genes were largely enriched for ribostatic and proteostatic functions (**Figure 13B, Bottom**). Next, given that these genes were identified through a co-expression analysis, we sought to identify potential regulatory logic to understand co-regulation of these driver genes. Using the ChEA and ENCODE databases, we assessed enrichment for transcriptional regulators that could regulate the driver genes. As with the thresholded upregulated genes (**Table 5-4**) we once again observed a signature of regulation by PRC2 and REST along with GATA and CTCF for the mouseUP driver genes (**Figure 13C, Left**). The mouseDOWN driver genes were enriched for TAF1 regulated genes, and genes regulated by BRCA1, YY1, MAX, and E2F4 (**Figure 13C, Right**). Intriguingly, within the E2F4-regulated gene list were two core components of the PRC2 – *Suz12* and *Eed*, which were downregulated in V_321_L DG compared to WT DG.

Genes clustered within a WGCNA module have been shown to be functionally related (Kos, Duan et al. 2018). Thus, we asked whether the genes with correlated expression in each module encode proteins that have functional interactions. We accomplished this using the STRING protein interaction (PI) database (Szklarczyk, Kirsch et al. 2023) to build PI Networks (PINs) from the WGCNA modules. Using stringent criteria (see methods) we extracted a PI network using genes within the WGCNA modules (**Figure 14**). We found that the protein products of genes within the WGCNA modules had highly significant functional interactions forming densely interconnected networks (**Figure 14A, mouseUP module**: 1136 nodes, average node degree=8.2, PI enrichment p value= 3.56e-13; **Figure 14C, mouseDOWN module**: 712 nodes, average node degree=15, PI enrichment p value < 1.0e-16.).

**Figure 14:**
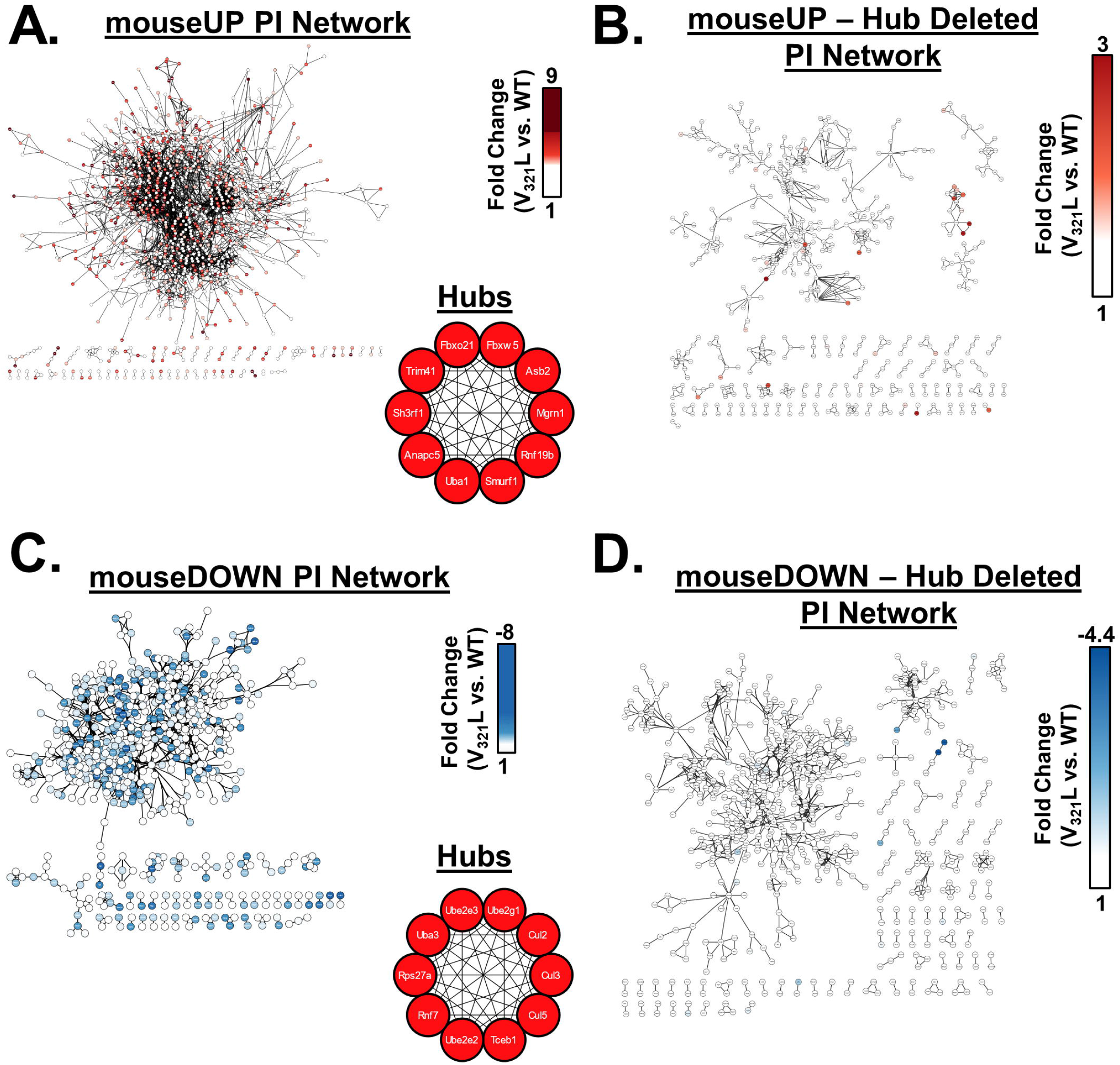
Ubiquitin ligases maintain network cohesion in protein interaction networks of V_321_L mouse WGCNA modules. Protein interaction (PI) networks were constructed (See methods for details) from WGCNA modules. Resulting networks were densely interconnected. Nodes are color coded according to fold change in expression (V_321_L vs. WT) indicated by the heatmaps in the respective graphs. Edges were bundled for clarity. **(A) mouseUP PI Network:** PI network from genes positively correlated (upregulated) with the V_321_L genotype (1910 nodes, average node degree=3.2, PPI enrichment p value= 3.56e-13). Node color legend: fold-change 1 white to 9 deep red. (**Inset**) mouseUP PI network was analyzed using MCC to find a ranked list of hub proteins within the network. The top 10 hubs for this network were all co-ranked as a rank 1 (red) and were ubiquitin ligases or accessory factors. **(B)** Hub genes embed the top upregulated genes in the V_321_L DG in the mouseUP PI network. The panel shows a PI network from genes positively correlated (upregulated) with the V_321_L genotype after the MCC hub genes were removed from the network and disconnected nodes were removed (479 nodes, average node degree=3.14). Node color legend: fold-change 1 white to 3 deep red. Also see **Extended Data Table 14-1**. **(C) mouseDOWN PI Network:** PI network from genes negatively correlated (downregulated) with the V_321_L genotype (1870 nodes, average node degree=4.7, PI enrichment p value < 1.0e-16). Node color legend: fold-change -8 blue to 1 white. (**Inset**) mouseDOWN PI network was analyzed using MCC to find a ranked list of hub proteins within the network. The top 10 hubs for this network were all co-ranked as a rank 1 (red) and were ubiquitin ligases or accessory factors. **(D)** Hub genes embed the most downregulated genes in the V_321_L DG in the mouseDOWN PI network. The panel shows a PI network from genes negatively correlated (downregulated) with the V_321_L genotype after the MCC hub genes were removed from the network and disconnected nodes were removed (577 nodes, average node degree=4.38). Node color legend: fold-change -4.4 blue to 1 white. Also see **Extended Data Table 14-1.**

To gain insight into the nature of these functional interactions, we analyzed the networks as undirected graphs. We identified hub nodes based on their maximal clique centrality (MCC) (Chin, Chen et al. 2014). The highest MCC ranked genes in the mouseUP and mouseDOWN modules were members of ubiquitin ligase complexes indicating that a functionally related group of hub genes was disrupted within these modules (**Figure 14A and C insets**). To probe the significance of these hub genes in maintaining integrity of the larger network and validate their hub status, we removed them from the network and reanalyzed the remaining genes for interactions in STRING for both mouseUP and mouseDOWN modules. Deletion of the MCC hub genes resulted in reduced connectivity within the network and disintegration of the network (**Table 14-1**) thereby supporting their role as hub genes; the hub deleted network resulted in loss of almost all the highest differentially expressed genes between V_321_L and WT DG as evidenced by the loss of darker colored nodes from the networks (**Figure 14 B&D**).

Thus, WGCNA revealed that loss of Nrg1 nuclear back signaling results in highly correlated dysregulation of functionally related modules of genes.

### Molecular changes in the mouse V_321_L mutant DG show similarity with those observed in DG samples from humans with schizophrenia

The bulk of the genetic variation associated with schizophrenia is regulatory in nature (eQTLs) (Ripke, Neale et al. 2014). However, available databases do not yet provide clear mechanisms for how these variants might influence biological processes that underlie schizophrenia pathology. Our network analyses and ontology analyses on the V_321_L mouse DG transcriptome allowed us to uncover latent regulatory structure within the transcriptomic data. We next sought to apply the same network analysis pipeline to anatomically matched human DG transcriptomic data from people with schizophrenia to ask which, if any, features uncovered in the analysis of the V_321_L mouse DG might be represented in human pathology.

We performed WGCNA using published RNASeq data from microdissected DG granule cell layer from healthy controls and people with schizophrenia (93 controls and 75 schizophrenia subjects: 119 males and 49 females) (Jaffe, Hoeppner et al. 2020) (**Figure 15**). WGCNA identified 2 modules wherein gene expression was significantly (positively or negatively) correlated with schizophrenia diagnosis (**Figure 15 A&B**). We renamed the module colors for clarity to parallel the data from the mutant mouse DG (Blue module - humanUP module; p=0.0004, corr.coefficient (Schizophrenia)= 0.3, corr.coefficient p=0.0005) (Turquoise module-humanDOWN module; p=0.0002, corr.coefficient (Schizophrenia)= -0.3, p=0.0005) and not with sequencing batch, race or sex of the individuals (**Figure 15B**). We also performed permutation analysis to validate the discovered modules as we did for the mouse DG samples and did not find values that matched or exceeded the actual values (**Figure 15C**).

**Figure 15:**
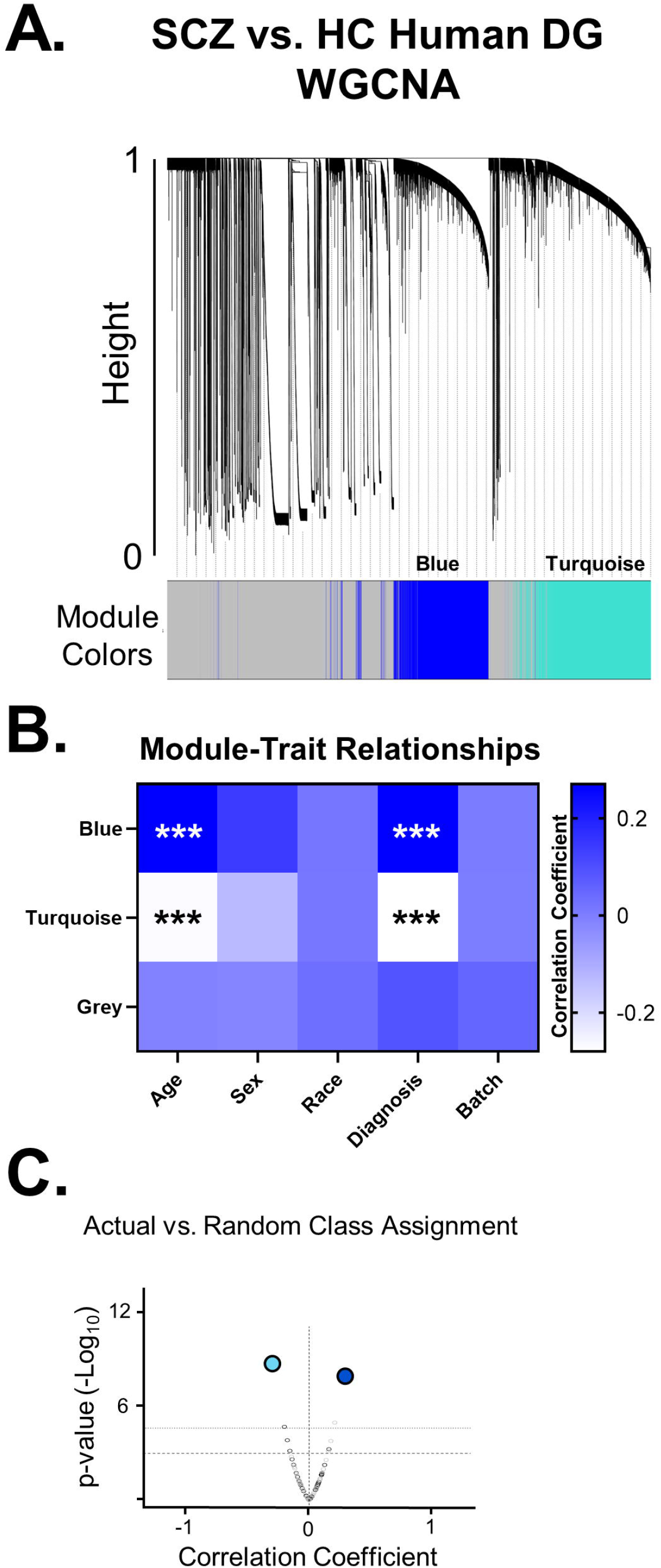
Weighted Gene Correlation Network Analysis (WGCNA) of human schizophrenia DG transcriptomes. **(A)** Module detection (WGCNA) via dynamic tree cut from the healthy control vs. schizophrenia DG transcriptome. The detected modules are represented by colors below the dendrogram. The two statistically significant modules (Turquoise and Blue) are labeled. **(B)** Heatmap showing module-trait relationships between detected modules in panel A and schizophrenia diagnosis, biological factors such as age and sex of the subjects, and race of the subjects and sequencing batch. The color indicates the correlation coefficient according to the legend. Only two modules were statistically significantly associated with diagnosis and age but not with sequencing batch or race. We dubbed the Turquoise module as the **humanDown module** (module x diagnosis: p=0.0002; module x age: p=0.0005) and the Blue module as the **humanUp module** (module x diagnosis: p=0.0004; module x age: 0.0005) for a more intuitive nomenclature. **(C)** Permutation analysis of module discovery-gene memberships were permuted 40 times. None of the permuted modules had equivalent correlation coefficients and statistical significance as the observed modules. The grey open circles represent permuted Turquoise modules and black open circles represent permuted Blue modules.

To validate that the human WGCNA modules represented functionally interacting components we constructed PINs from them (**Figure 16A&C**). We found that the protein products of genes that were part of the human WGCNA modules had highly significant functional interactions (**Figure 16A, humanUP module**: 489 nodes, average node degree= 8.96, PI enrichment p value= 1.96e-7. **Figure 16C, humanDOWN module**: 1180 nodes, average node degree= 23.56, PI enrichment p value= <1.0e-16). Similar to the mouse hub gene analysis, the highest ranked hub genes formed networks that are part of ubiquitin ligase complexes (**Figure 16A&C insets**). Deletion of the human MCC hub genes resulted in loss of almost all the relatively differentially expressed genes between SCZ and HC groups as evidenced by the loss of darker colored nodes from the networks (**Figure 16 B&D**).

**Figure 16:**
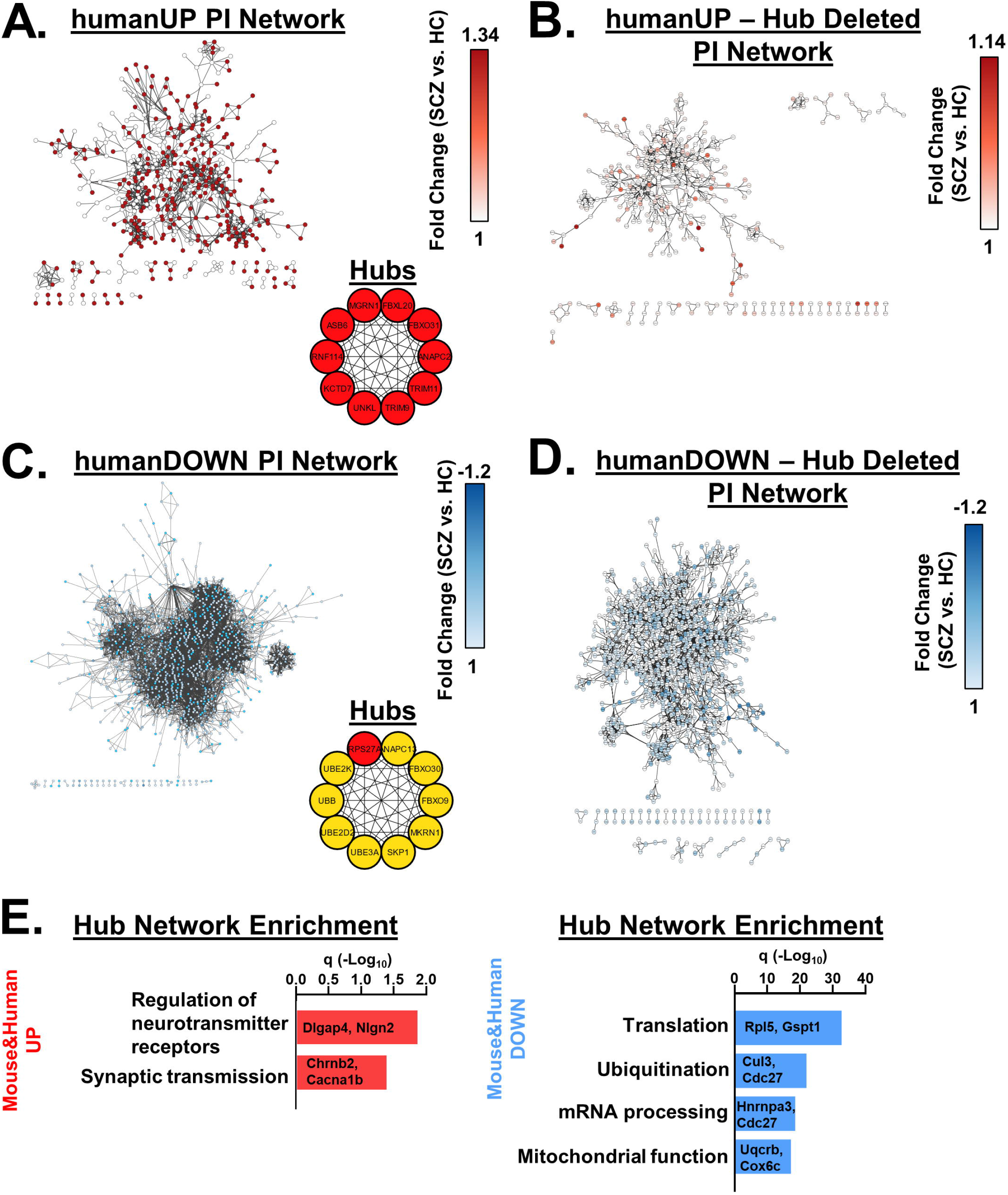
Shared functional molecular risk profile between V_321_L mutant mouse DG and postmortem DG samples of people who were diagnosed with schizophrenia. **(A) humanUP PI Network:** PI network from genes positively correlated (upregulated) with schizophrenia diagnosis (1364 nodes, average node degree= 8.96, PI enrichment p value= 1.96e-07). Node color legend: fold-change 1 white to 1.14 deep red. (**Inset**) humanUP PI network was analyzed using MCC to find a ranked list of hub proteins within the network. The top 10 hubs for this network were all co-ranked as a rank 1 (red) and were ubiquitin ligases or accessory factors. **(B)** Hub genes embed the top upregulated genes in the DG of patient samples in the humanUP PI network. The panel shows a PI network from genes positively correlated (upregulated) with the schizophrenia diagnosis after the MCC hub genes were removed from the network and disconnected nodes were removed (361 nodes, average node degree=7.9). Node color legend: fold-change 1 white to 1.14 deep red. **(C) humanDOWN PI Network:** PI network from genes negatively correlated (downregulated) with schizophrenia diagnosis (1947 nodes, average node degree= 23.56, PI enrichment p value= <1.0e-16). Node color legend: fold change -1.2 blue to 1 white. (**Inset**) humanDOWN PI network was analyzed using MCC to find a ranked list of hub proteins within the network. The network had one rank 1 hub gene (*RPS27A*-red) and the other 9 were co-ranked for rank 10 (yellow). All hub genes were identified as ubiquitin ligases or accessory factors. **(D)** Hub genes embed the top downregulated genes in the DG of patient samples in the humanDOWN PI network. The panel shows a PI network from genes negatively correlated (downregulated) with schizophrenia diagnosis after the MCC hub genes were removed from the network and disconnected nodes were removed (1064 nodes, average node degree=22.47). Node color legend: fold change -1.2 blue to 1 white. **(E)** Gene Ontology analysis of the up and downregulated genes that comprised the hub genes identified in both V_321_L mouse and SCZ human PINs and their first-degree neighbors from their respective networks showed complete overlap of the biological processes predicted to be altered. Examples of common differentially expressed genes between species are displayed.

We created hub networks by selecting the first-degree neighbors for the mouse and human hub genes and asked what biological processes were enriched in these core networks. We found a striking 100% overlap between the mouse and human hub network functions. The mouse and human UP hub networks were enriched for genes involved in synaptic transmission and neurotransmitter receptor regulation by synaptic proteins (**Figure 16E, Left**). The mouse and human DOWN hub networks were enriched for genes involved in several cellular homeostatic functions such as proteostasis, ribostasis, and mitostasis (**Figure 16E, Right**).

Thus, these data reveal evolutionarily conserved regulatory programs that might regulate expression and/or function of genes that underlie cellular pathology in psychotic disorders. Furthermore, these data highlight the potential for convergence of molecular pathology of rare and common genetic variants associated with schizophrenia.

## DISCUSSION

We investigated the cellular and molecular consequences of loss of Nrg1 nuclear back signaling. Loss of nuclear back signaling by the Nrg1 ICD in the V_321_L mouse results in transcriptional changes in the DG and alterations to cell cycle dynamics, neurogenesis, neuronal maturation, and schizophrenia-related gene expression. These changes were accompanied by pronounced sensorimotor gating deficits. Network analyses uncovered latent gene regulatory factors and functional logic of gene co-expression networks dysregulated by the mutation. Comparison of the V_321_L mouse DG transcriptome with human schizophrenia DG transcriptomes using WGCNA revealed ubiquitin ligases as hubs coordinating large protein interaction networks involved in synaptic transmission, RNA processing/trafficking and mitochondrial function.

### Balance between back signaling mechanisms: a tight rope for circuit development

Type III Nrg1 KO neurons show stunted growth of axons and dendrites, which was rescued by re-expression of the full length Type III Nrg1 protein (Chen, Hancock et al. 2010). The Type III isoform of Nrg1 is unique in that it localizes to the axonal pre-synapse indicating that the effects of Nrg1 signaling on axon and dendrite growth require axon localized Nrg1 protein (Vullhorst, Ahmad et al. 2017). Axon and dendrite outgrowth are regulated by two distinct Nrg1 functions: Nrg1-dependent axonal growth requires activation of PI3K signaling (local back signaling) (**Figure 2D**) whereas dendrite growth requires nuclear back signaling (**Figure 2E**). It is likely that these although these two signaling functions appear to be independent, they are coordinated. PI3K signaling is a membrane associated signaling pathway, and membrane fractions from V_321_L brains showed higher levels of Nrg1 protein (**Figure 2B**). It is possible that increased membrane residence time of the V_321_L Nrg1 ICD resulting from decreased proteolysis, increases the duration of PI3K signaling and enhances local signaling. Indeed, the data in **Figure 2D (as well as in (Chen, Hancock et al. 2010))** show trends towards longer axon lengths in V_321_L neurons, and treating granule cell cultures with the gamma secretase inhibitor increased the number of primary neurites that were immunoreactive for axonal neurofilaments (**Figure 4B, 4D**). Based on these findings we propose that activation of PI3K in Type III Nrg1 expressing axons supports axonal trafficking as previously shown (Hancock, Canetta et al. 2008, Zhong, Akmentin et al. 2022). The same stimuli capable of activating local back signaling can also activate gamma secretase cleavage of axonal Nrg1 initiating nuclear back signaling. The nuclear ICD downregulates genes related to early axon growth programs and simultaneous upregulates dendritic maturation gene programs (**Table 6-2**). In this model, the nuclear ICD functions to couple axonal growth and target innervation to dendritic arborization. It is well documented in cultured hippocampal neurons and as well as in developing granule cells *in vivo*, that axonal growth precedes dendritic maturation (Dotti, Sullivan and Banker 1988, Sun, Sailor et al. 2013). We propose that the balance between local and nuclear Nrg1 back signaling mechanisms represents a potential channel for afferent and efferent synaptogenesis to be co-ordinated within a neuron.

### How does the Nrg1 ICD regulate gene expression?

The Nrg1 ICD lacks any obvious DNA binding domains; however, it has strong transactivation properties, rivaling those of the VP16 transactivation domain (Bao, Wolpowitz et al. 2003). The upregulated genes in the V_321_L DG were predicted to be targets of the PRC2, the catalytic subunits (*Eed, Ezh2* and *Suz12*) of which were downregulated in the V_321_L DG. How Nrg1 nuclear signaling regulates the expression and/or function of the PRC2 is not known but antagonistic interactions between PRC and the SWI/SNF chromatin remodeling complexes have been noted in the context of maintaining balance between self-renewal and differentiation of stem cells (Seo and Kroll 2006, Kadoch, Copeland and Keilhack 2016, Weber, Hafner et al. 2021). Interestingly, the Nrg1 ICD was shown to interact with Brm, a core ATPase subunit of the SWI/SNF complex and BAF57 another subunit of the SWI/SNF complex in cultured neural stem cells (Pirotte, Wislet-Gendebien et al. 2010). It is possible that the Nrg1 ICD regulates competition between these large remodeling complexes to influence cell fate decisions.

We also noted downregulation of several ribosomal subunit genes and *Cenp* (centromere proteins that couple centromeric chromatin to the kinetochore) genes in the V_321_L DG (**Tables 5-1**). The kinetochore genes and genes related to regulation of microtubule dynamics are predicted to be E2F4 target genes (**Table 5-5**). The E2F family of transcription factors can function as both activators and repressors. E2F4 is one of three canonical E2F repressors along with E2F5 and E2F6 (Chen, Tsai and Leone 2009). This implies a potential gain of function of E2F4 in the V_321_L mutant. Repressor E2Fs have been shown to interact with the PRC and E2F4 accumulates in quiescent cells consistent with its role as a factor that maintains differentiated states (Trimarchi, Fairchild et al. 2001, Cuitiño, Pécot et al. 2019). Thus, it is possible that both, the upregulated and downregulated genes in the V_321_L mice result from disrupting PRC occupancy across the genome.

Our findings suggest that PRCs might be temporally regulated during neuronal development to target various genes associated with a common function. In line with this, the presence of “polycomb domains”, which are clusters of polycomb proteins bound to DNA segments located several megabases from each other but located in the same 3D space have been demonstrated (Blackledge and Klose 2021). These domains are coordinated by CTCF-cohesin complexes (Rhodes, Feldmann et al. 2020). Genes dysregulated in the V_321_L mutant that are predicted to be regulated by CTCF-cohesin shared functional overlap with the predicted PRC2 target genes (**Tables 5-4 and 5-5**).

### Nrg1 nuclear back signaling and genetic risk for psychosis

Genetic variants that influence gene expression, eQTLs, are thought to comprise a substantial proportion of genetic risk in schizophrenia (Richards, Jones et al. 2012). The cellular mechanisms by which these changes in gene expression influence risk for schizophrenia are not clear. We found a significant enrichment of genes associated with psychotic disorders to be dysregulated in the V_321_L DG (**Tables 5-3 and 11-1**). We examined known interaction networks between these genes and found several modular hubs with hub genes represented by the known schizophrenia-associated genes. This larger network of genes was enriched for transcriptional regulators with known regulatory variants associated with schizophrenia (**Figure 11**). These data are in line with predictions from the omnigenic model wherein core genes/functions are part of densely connected cellular networks of peripheral genes/functions (Boyle, Li and Pritchard 2017). Additionally, since the core genes involved in synaptic transmission can be accessed by many peripheral genes with cellular homeostatic functions, our data also inform potential mechanisms explaining how peripheral genes might have a cumulative effect on heritability of schizophrenia. Intriguingly, while hub genes identified in our SCZ+ network are not obvious mediators of synaptic function (**Figure 11**), half of them have been shown to be part of the hippocampal synaptic proteome in both rodents and humans (Koopmans, Pandya et al. 2018). Strikingly, we observed a very similar profile and high functional overlap when we performed additional network analyses on a published dataset of human schizophrenia and healthy control DG granule cell layer transcriptomes (Jaffe, Hoeppner et al. 2020). In both V_321_L mouse and schizophrenia human networks, ubiquitin-proteosome system (UPS) associated genes formed the hubs (**Figures 14 and 16**). The high degree of functional overlap between the genetic dysregulation of the V_321_L mouse and human schizophrenia DG samples indicates that there might indeed be convergence between risk factors at the gene regulatory level since it is unlikely that many of the patients whose DG RNASeq data were used were carrying the V_321_L mutation in Nrg1 (∼ 0.2% of the people would be predicted to be homozygous for this mutation). Future studies examining gene regulatory network overlaps between various genetic and environmental models of schizophrenia risk could test for such convergence.

### Concluding remarks

Psychiatric-disease risk variants across various disorders aggregate in signaling pathways involved in protein modification such as the UPS and histone modifications (O’dushlaine, Rossin et al. 2015). We found that that disease-associated pathways can be regulated by a common set of transcriptional regulators such as the PRCs. We propose that these regulators can in turn be regulated by specific neurodevelopmental signaling networks. The combination of transcriptional regulators impinging upon gene networks within “risk-associated pathways” and intersecting developmentally regulated signaling events might provide specificity for specific disorders and offer a framework to understand how environmental factors interact with genetic risk associated with disease.

## METHODS & MATERIALS

### SUPPLEMENTAL MATERIALS AND METHODS

#### KEY RESOURCES TABLE

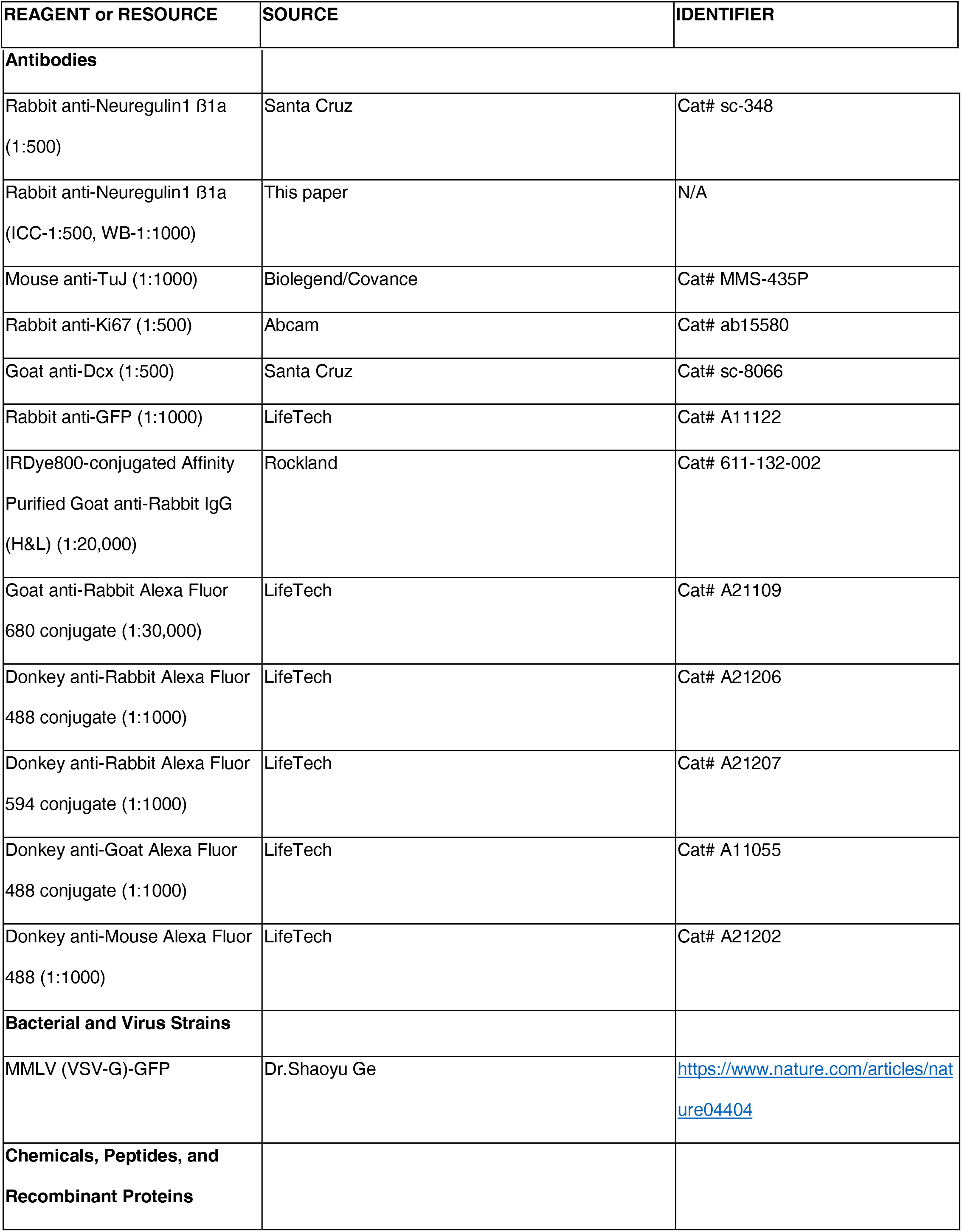
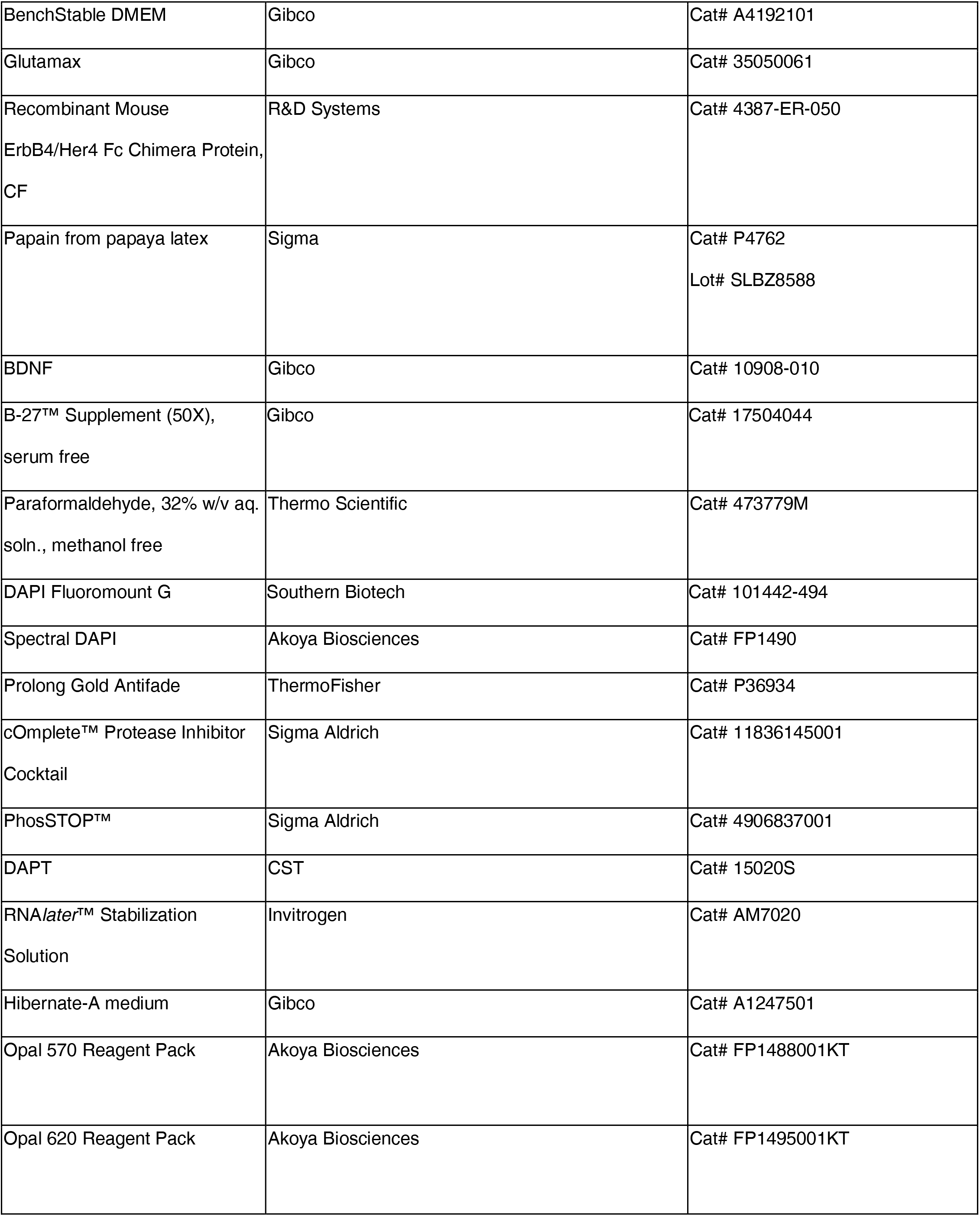
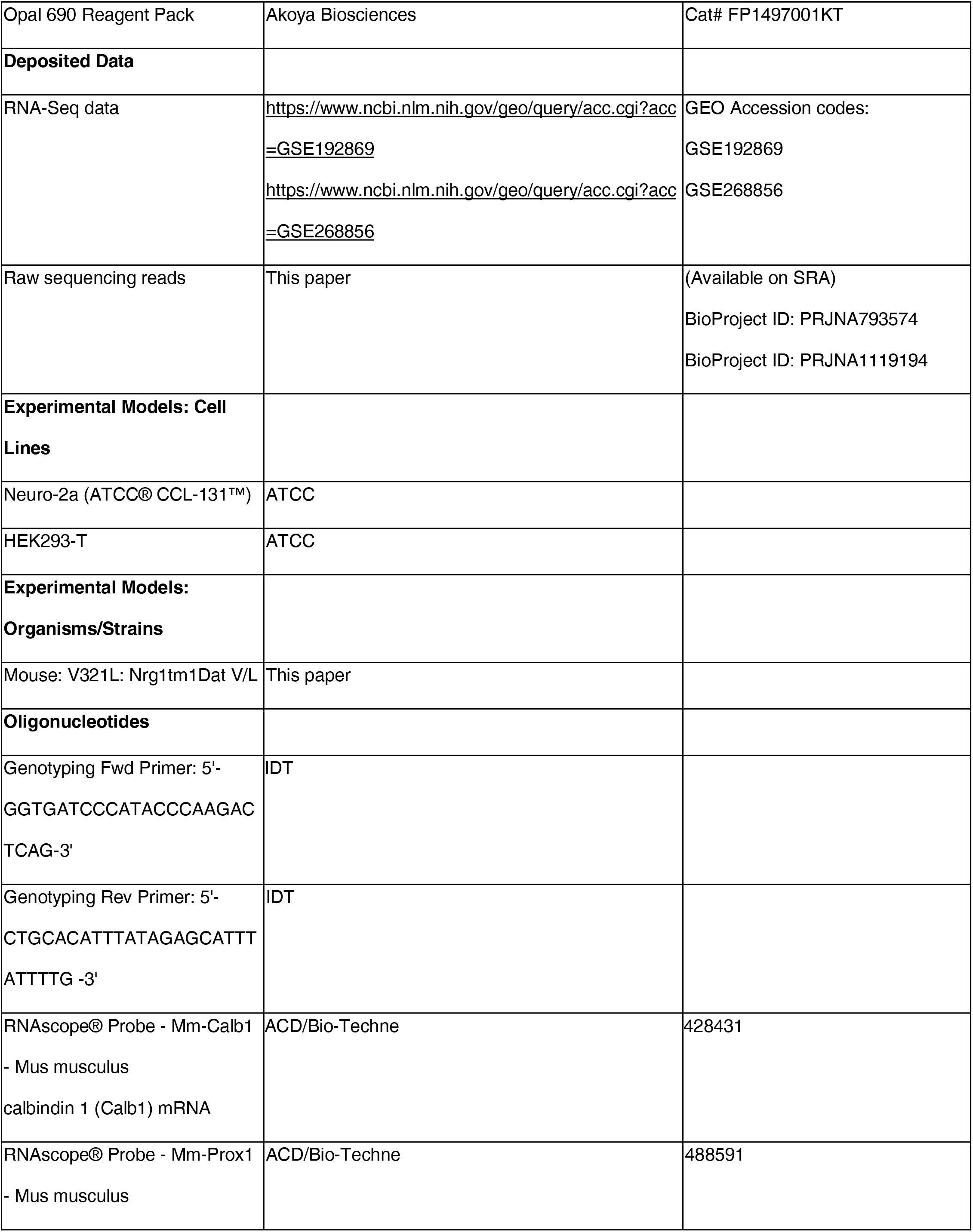
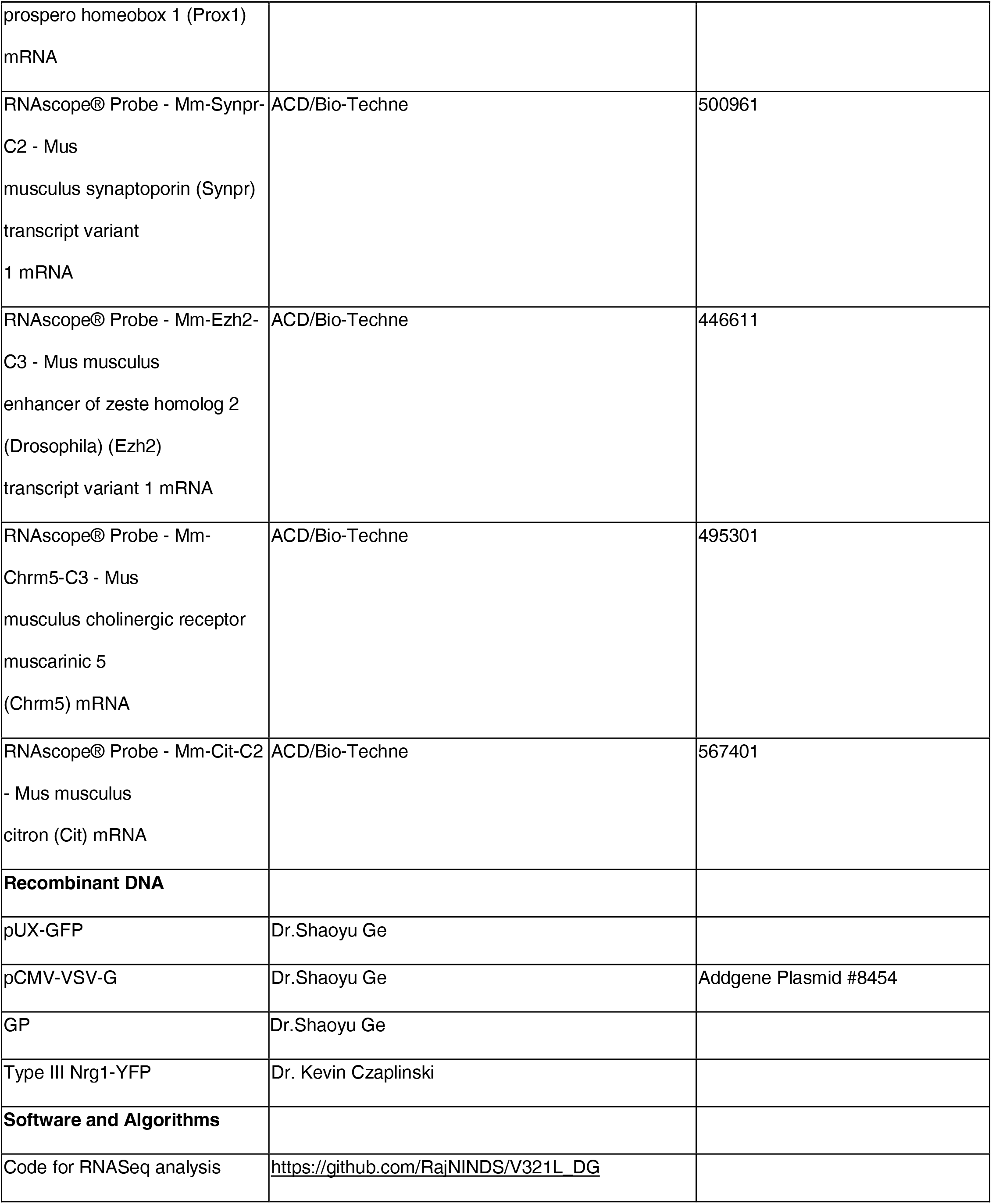

##### Experimental model and subject details

Male and female mice ages E18.5 - 6 months were used. Animals were housed in a 12-hour light/dark cycle (lights on from 7 AM to 7 PM), environment that was both temperature and humidity controlled. Animals had free access to food and water. All animal care and experimental procedures were approved by the Institutional Animal Care and Use Committee of the SUNY Research Foundation at Stony Brook University, Stony Brook NY (Protocol #1618) and NINDS, National Institutes of Health, Bethesda MD (Protocol #1490).

For studies profiling proliferating cells in the DG, mice around the age of P21 were used to avoid a “floor effect” due to decline in proliferating cells with age (Ben Abdallah, Slomianka et al. 2010). Mice aged ∼2 mo. were used for counting Dcx+ cells as Dcx immunoreactivity is still fairly abundant in the DG at this age. For studies of cell cycle dynamics, we used mice around the age of P90 as death of proliferating cells is high in younger mice and stabilizes around this age thereby allowing us to detect the proliferative state of the same cell using EdU and Ki67 (Ansorg, Witte and Urbach 2012). For the morphological studies we intended to profile enduring changes in mature GCs. The bulk of granule cells undergo a relatively “sudden” maturation (profiled by gene expression changes) around the 3^rd^ postnatal week (Hochgerner, Zeisel et al. 2018). Further, complete morphological maturation of a GC takes ∼1 month (Sun, Sailor et al. 2013). Thus, to escape the dynamic period of early maturation and presence of large numbers of immature neurons in the granule cell layer we assessed morphology of GCs after 2 months of age.

##### Genotyping

Genotypes were determined by PCR using the following primers:

> NDEL1 - Forward primer 5’- GGTGATCCCATACCCAAGACTCAG -3’

> NDEL2-Reverse primer 5’- CTGCACATTTATAGAGCATTTATTTTG -3’

##### Pre-pulse inhibition

Male and female mice between the ages of 1-3 months were used. Testing was performed in dark sound attenuated boxes, between the hours of 12pm and 4pm; male and female mice were tested separately, and all equipment was thoroughly cleaned between mice using Clorox Fuzion® cleaner allowing ∼2 min contact time with the equipment. All testing was performed using the SR-LAB-Startle Response System (San Diego Instruments). Mice were transported to the behavior testing facility and held in a holding room. Mice were acclimated to the holding room for at least 5 min and then to the startle chambers by placing them into the startle chamber and exposing them to constant background white noise set at 65dB for 5 min for three days. On the third day, mice also underwent an input-output curve calibration where they were exposed to startle stimuli increasing in 5dB increments from 70dB to 120dB, an input-output curve was plotted for each mouse to identify saturation of the startle response. All mice tested showed maximal startle responses at 115dB which was used as the startle stimulus in the Habituation and PPI trials. On the fourth day mice were once again exposed to 65dB background white noise for 5 min after which they entered “Block 1” consisting of sixty 20ms 115dB startle stimuli with an intertrial interval of 20s. Block 1 continued into “Block 2” consisting of ten trials each of the following combinations – 4ms 68/70/75/85dB pre-pulses followed by a 115dB startle stimulus separated by an interval of either 30 or 100ms. Twenty 20ms startle stimuli (115dB) were interspersed between these trials. Intertrial intervals were randomized using the ITI function within the SR-LAB software and the trial order was randomized using an atmospheric noise randomizer (www.random.org/lists). PPI is known to improve upon repeat testing, stabilizing thereafter (Valsamis and Schmid 2011). Thus, on the fifth day, mice were once again exposed to Blocks 1 and 2. Responses from day 5 were used as the outcome measures. Additionally, fecal pellets were counted after every session and recorded for each mouse.

###### PPI calculation

Startle responses were identified as the maximal voltage deflection recorded from the startle box during the 20ms startle pulse delivery. An average of the responses to the twenty startle stimuli alone delivered in Block 2 were used as the baseline startle measurement to calculate %PPI. %PPI was calculated as [(Average startle response during PPI trial/baseline startle response) *100]. All %PPI values were capped at a lower limit of 0% (thus any negative PPI was set to and reported as 0% PPI).

##### Antibody generation

Antibodies recognizing the Nrg1 intracellular domain were raised in rabbits using the immunizing peptide: DEE(pY)ETTQEYEPAQEP by Biomatik. Antibodies recognizing the phosphorylated peptide DEE(pY)ETTQEYEPAQEP vs. the unphosphorylated peptide DEEYETTQEYEPAQEP were purified by affinity chromatography. A cocktail of antibodies referred to as anti-Nrg1 ICD_DAT_ was used for staining cells comprised of antibodies against the phosphorylated and non-phosphorylated peptides. The phospho-specific antibodies were developed for a different study. Antibodies were validated for Immunofluorescence and immunoblotting by comparing samples from N2A cells and N2A cells transfected with a Type III Nrg1-YFP construct. Detectable staining of N2A cells in ICC experiments and unique bands corresponding to the appropriate molecular weight was only observed in transfected samples.

##### Subcellular fractionation

For each experiment, cortex, and hippocampus were pooled by genotype for nuclear fractionation. P86-108 animals were deeply anesthetized with isofluorane and decapitated, and cortex and hippocampus were isolated on ice and homogenized with Dounce homogenizer on ice in 5mL nuclear fractionation buffer (60mM KCl, 15mM NaCl, 15mM Tris-HCl pH7.6, 1mM EDTA, 0.2mM EGTA, 15mM 2-mercaptoethanol, 0.5mM spermidine, 0.15 spermine, 0.32M sucrose, cOmplete ULTRA tablets protease inhibitor cocktail). Samples were first homogenized with the loose pestle checking for single, dispersed cells after every 10 strokes using a hemocytometer. Samples were then homogenized with the tight pestle checking for dissociated nuclei after every 10 strokes. Samples were centrifuged at 1000 *xg* for 10 minutes at 4°C. The supernatant (S1) was preserved for further processing. The pellets were resuspended in nuclear fractionation buffer with 0.1% Triton X-100 and incubated on ice for 10 minutes and then pelleted at 1000 *xg* for 10 minutes at 4°C. Pellets were resuspended in 500μL nuclear fractionation buffer, layered over 500μL of a 1M sucrose (in nuclear fractionation buffer) cushion, and centrifuged at 1000 *xg* for 10 minutes at 4°C. Pellets were resuspended in 1mL nuclear fractionation buffer.

To isolate the membrane fractions, fraction S1 was centrifuged at 10,000 xg for 10 minutes at 4°C to remove mitochondria, lysosomes, and peroxisomes. The supernatant (S2) was collected and centrifuged at 100,000 xg for 1 hour at 4°C. The pellet was resuspended in 1mL subcellular fractionation buffer (SFB) (250mM Sucrose, 20mM HEPES, 10mM KCl, 1.5mM MgCl_2_, 1mM EDTA, 1mM DTT (add fresh before use), cOmplete ULTRA tablets protease inhibitor cocktail) and passed through a 25-gauge needle 10 times, and the sample was centrifuged at 100,000 xg for 45 minutes at 4°C. The supernatant was discarded, and the pellet was resuspended in 200μL SFB.

Laemmli buffer was added to each sample and boiled at 95°C for 5 minutes. Denatured samples were stored overnight at -20°C. Protein analysis was then performed by immunoblot.

##### Immunoblotting

Samples were electrophoresed on 8% SDS-PAGE gel at 90V for 1.5 hours at room temperature. Proteins were transferred onto nitrocellulose membranes via wet transfer performed at 4°C at 100V for 1 hour. Membranes were blocked in Odyssey Blocking Buffer (LI-COR #927-40000) for 1 hour at room temperature with gentle agitation. Membranes were then incubated overnight on a rocker at 4°C with primary antibodies diluted in 50% blocking buffer. The next day, membranes were washed 3 times with PBS-T (PBS+0.1% Tween20) and incubated with secondary antibodies (diluted in 50% blocking buffer) for 1 hour at room temperature with gentle agitation. After 3 washes in PBS-T, membranes were imaged using the LI-COR Odyssey Infrared Imaging System.

##### Cell culture

###### Primary neuronal culture

Postnatal day 4-5 WT and V_321_L mice were rapidly decapitated, and their brains were transferred to ice-cold Hibernate-A. Hippocampi were dissected out and transferred to Hibernate-A (ThermoFisher) on ice. Hibernate solution was removed and replaced with activated and filtered Papain (Sigma Lot# SLBZ8588) dissolved in Hibernate-A. Hippocampi were digested for 15min in a 37°C water bath gently inverting the tube every 5min. Papain solution was removed, and digested tissue was washed thoroughly (4x) with wash solution (Hibernate-A+10%FBS+100U DNase I (StemCellTech)) by resuspending the tissue in wash solution and allowing it to settle before replacing wash solution. 40µm cell strainers were placed over 50mL conical tubes and wet with plating medium (Neurobasal+B27+Glutamax+Pen/Strep+HEPES) and flowthrough was discarded. Wide and narrow bore fire-polished Pasteur pipettes were pre-wet with FBS and were used to triturate the tissue into a single-cell suspension. The cell-suspension was applied to the pre-wet cell strainer and flow through was collected and analyzed for live/dead cells using Trypan blue on the CountessII automated cell counter. ∼100kcells/well were plated into 24 well plates containing 12mm Poly-D-Lysine+Laminin coated coverslips in a total volume of 250µL/well. Neurons were left overnight to adhere, and 250µL fresh plating medium was added the following day to each well. 50% of the total media was replaced every other day until experimental time points. For cortical cultures, cortices were isolated from E18.5 embryos.

###### Cell line culture and transfection

Neuro2A cells (ATCC) were maintained in DMEM+10%FBS+1:1000 Gentamycin. Cells were passaged on a Mon-Wed-Fri scheduled and were not used beyond 10 passages. Cells were transfected with plasmid DNA using Lipofectamine2000 following manufacturer guidelines. Plasmid details can be found in the Key Resources Table.

##### Stimulation & drug treatments

Recombinant Mouse ErbB4/HER4 Fc Chimera Protein was reconstituted at a final concentration of 0.5µM and stored at -80°C. Working stocks of 0.1µM were prepared and stored briefly at -20°C. For stimulation experiments, 20nM ErbB4 solution was prepared by diluting 0.1µM stock solution in 125µL Neurobasal plating medium/well.

Neurons were washed by gently swirling the plate and aspirating out all the media and near-simultaneously replacing with 125µL of fresh pre-warmed plating media and equal volume of Drug concentrations, (pre)treatment periods are listed in the table below:

| Drug Name | Manufacturer | Concentration | Treatment Time |
| --- | --- | --- | --- |
| DAPT | CST | 20μM | 24h, 45min |
| sErbB4 | R&D Systems | 20nM | 15-20min |
| Wortmannin | Sigma | 500nM | 45min |
| L685458 | Sigma | 5mM | 45min |

##### Immunocytochemistry

Neurons were fixed for 5min at room temperature (RT) in 4% PFA by adding equal volume of 8%PFA to neurobasal in the wells. Entire solution was replaced with fresh 4%PFA and neurons were fixed further for 10min at RT. Coverslips were washed 3x 5min each in 1xPBS-G (PBS+0.1M Glycine), incubating in the final wash for 10min. Neurons were then permeabilized in 0.1% Triton X-100 in 1xPBS for 15min at RT. Coverslips were washed 3x 5min each with 1x PBS-G and incubated in blocking solution (10% normal donkey serum-NDS) for 30min. After blocking, coverslips were incubated with primary antibodies prepared in 1% NDS overnight at 4°C. Coverslips were washed thoroughly with 1xPBS at RT 3x 5min each and were then incubated with secondary antibodies prepared in 1%NDS+0.1%Triton X-100. Coverslips were washed thoroughly with 1xPBS at RT 3x 5min each and mounted in mounting medium containing DAPI (DAPI Fluoromount-G™ - Southern Biotech). Antibodies used and concentrations can be found in the Key Resources Table (KRT).

##### Tissue processing and immunohistochemistry

Animals were deeply anesthetized with isofluorane and transcardially perfused with 4% paraformaldehyde (PFA) in PBS. Brains were harvested and post-fixed overnight at 4°C in 4% PFA in PBS. Brains were transferred to 30% sucrose in PBS and incubated at 4°C with agitation until they sank. Brains were embedded in optimal cutting temperature (O.C.T.) compound, flash frozen and stored at -80°C. 20-50μm sections were obtained serially over the entire dorsoventral extent of the hippocampus, and each slide collected 1 of every 8 (20 μm) or 1 of every 4 sections (50 μm). Sections were dried at room temperature overnight prior to immunohistochemistry.

Antigen retrieval was performed for Ki67 staining by immersing sections in antigen retrieval buffer (10mM sodium citrate, 0.05% Tween20, pH 6.0) at 100°C for 15 minutes. Sections were then allowed to cool to room temperature and neutralized in 0.1M borate buffer, pH8.5 for 10 minutes at room temperature. After 3 5-minute washes in PBS, sections were incubated in blocking buffer (5% normal donkey serum, 0.1% Triton X-100 in PBS) for 1 hour at room temperature. Sections were then incubated with primary antibodies diluted in blocking buffer for 48 hours at 4°C in a humidity chamber. After 3 5-minute washes in PBS, sections were then incubated with secondary antibodies diluted in blocking buffer for 1 hour at room temperature. Sections were dehydrated in 70% ethanol for 5 minutes and treated with autofluorescence eliminator reagent (Millipore Sigma 2160). No. 1.5 glass coverslips were then mounted on the slides with DAPI Fluoromount-G and allowed to dry overnight at room temperature.

Antibodies used and concentrations can be found in the Key Resources Table (KRT).

##### Fluorescent in-situ hybridization (FISH)

Brains were fixed and cryosectioned as described earlier. 15µm sections were prepared with two coronal sections on a slide. FISH was performed per manufacturer’s instructions using the RNAscope v2 fluorescent multiplex assay (ACD/Bio-Techne, CA). Details on the probes used to can be found in the KRT.

##### Imaging & image analysis

To image Nrg1 ICD puncta in the nucleus, imaging was done on the Zeiss LSM800 confocal microscope using the 63x oil immersion objective. All samples were imaged using identical parameters. Neurons to analyze were identified by expression of Type III Nrg1 (CRD+), and their nuclei were imaged at 2.5x digital zoom with 1µm z-steps.

Image processing and analysis was performed using ImageJ (FIJI). Briefly, center of the nucleus was identified and 1 z-section above and below it was flattened via maximum intensity projection. DAPI-stained nuclei were outlined, and the enclosed area was measured and noted. ICD clusters were manually counted using the cell-counter plugin. Density of clusters was calculated by dividing number of clusters by the nuclear area.

Images examining Type III Nrg1 expression profile in the dentate gyrus and images used for Ki67+ total cell counts in P21 animals were done using the Optical Fractionator probe on the Stereo Investigator software (MBF Bioscience) coupled to live imaging on an epifluorescent microscope. Section sampling fraction was 1 every 8th section, counting frame size used was 100 μm × 100 μm, and guard zone height used was 10 μm. All sample counts achieved a Gundersen coefficient of error of <0.1.

Dcx+ cell density counts were acquired on Olympus FLV1000 confocal microscope at 20X magnification with 3-μm z-steps. Images were analyzed in ImageJ using the Cell Counter plugin. Only cells located in the subgranular zone of the dentate gyrus were counted. Total cell counts for each animal were divided by the total dentate gyrus volume counted to obtain cell density. For each animal, 12-14 coronal sections were sampled along the entire dorsoventral extent of the dentate gyrus.

EdU labeling counts were done on images acquired using Olympus VS-120 microscope at 20X magnification. Images were acquired at 1-μm Z-steps of the entire dentate gyrus hemisection. Images were analyzed in ImageJ using the Cell Counter plugin. Total cell counts for each animal were divided by the total dentate gyrus volume counted to obtain cell density. For each animal, 12-14 coronal sections were sampled along the entire dorsoventral extent of the dentate gyrus.

RNAScope FISH sections were imaged on a Nikon Ti2 spinning disk confocal microscope using a 40x silicone immersion objective. Images were acquired at 1-μm Z-steps of the dentate gyrus. The images were Z-projected and an area in the molecular layer was identified to assess background staining. The fluorescence intensity of the background was measured and then subtracted using the Math>Subtract function on FIJI (ImageJ). Background subtracted images were then used to delineate an ROI consisting of the GCL using DAPI staining as a guide. The “IntDen” measurement within the GCL was recorded. 2 hippocampal sections (i.e. 4 hippocampi) at different anterior-posterior locations (consistent between mice) were quantified per mouse and averaged.

##### Golgi impregnation, sampling criteria, imaging, & analysis

P60 animals were deeply anesthetized with isofluorane, and brain samples were harvested quickly on ice and rinsed with ice-cold PBS. The brain samples were then treated with the FD Rapid GolgiStainTM Kit (FD NeuroTechnologies, Inc., catalog # PK401) per manufacturer’s guidelines. 100 μm brain sections were collected using a vibratome and dried overnight in the dark at room temperature. Sections were then dehydrated in progressively increasing concentrations of ethanol solution (50% 75% 95%), then xylene, and coverslipped with Permount solution.

Imaging of Golgi impregnated neurons in the dentate gyrus was done on the Olympus VS-120 microscope at 60x magnification. Individual neurons were selected for imaging based on the following **criteria**: 1) the neuron’s cell body resided in the granule cell layer in the intermediate section of the superior blade, 2) the soma morphology assumed the typical granule cell “tear drop” morphology, 3) the dendritic arbor originated from a single primary dendrite and assumed a cone-shaped morphology 4) the dendritic arbor was not obstructed by other cell bodies or dendrites. 11 neurons from 5 wildtype animals and 13 neurons from 5 homozygous mutants were ultimately selected for analysis. Dendritic arbor tracings, sholl analysis, and branching analysis were done on the NeuronStudio software.

##### GFP-expressing retrovirus production, injection, imaging, & analysis

GFP-retrovirus was prepared by co-transfection of pUX-GFP, VSVG, and GP constructs (kindly provided by the laboratory of Shaoyu Ge; previously described in (Ge, Goh et al. 2006)) into HEK293 cells. Viral supernatant was collected, the retrovirus was concentrated by ultracentrifugation, and the viral pellet resuspended in sterile PBS.

Following anesthetization, ∼P90 animals were stereotaxically injected bilaterally in dorsal dentate gyrus (-2.0 mm A/P, +1.6 mm M/L, -2.5 mm D/V) and ventral dentate gyrus (-3.0 mm A/P, +2.6 mm M/L, -3.2 mm D/V). The injection volume was 0.25 μL/site. Following surgery, animals were administered 0.03 mL ketorolac (3 mg/mL) for every 10g body weight.

14 days following stereotaxic viral injections, animals were sacrificed, and brain samples harvested as described previously for tissue preparation. 50-μm floating coronal sections were obtained, and each set of sections contained 1 of every 6th section of the hippocampus along its dorsoventral extent.

Analysis was done in real time while imaging on the Zeiss Axio Imager A1 Fluorescent Microscope using a 40x oil immersion objective lens. The identity of GFP+ cells were determined by a combination of morphology, Dcx immunoreactivity, and Nissl staining. Neural progenitors were identified by short, tangential processes and the lack of radially oriented processes, with or without Dcx immunoreactivity. Immature granule cells were identified by their radially oriented neurites, Dcx+ and Nissl+ staining. Astrocytes were identified by their star-like morphology, extensive branch ramifications, and lack of Dcx immunoreactivity and Nissl staining.

##### EdU labeling and tissue processing (cell cycle exit, length, and survival)

To measure cell cycle reentry, EdU (50 μg/g body weight) was injected intraperitoneally in animals at ∼P90. 24 hours after EdU injection, animals were sacrificed, and brain samples were harvested and prepared as described in the previous section. To visualize EdU incorporation, the Click-iTTM EdU Cell Proliferation Kit (Thermo Fisher Scientific, catalog # C10338) was used according to manufacturer’s guidelines, followed by immunohistochemistry as described in the previous section.

To measure cell cycle length, EdU (50 μg/g body weight) was injected intraperitoneally in animals at P21-30. 30 minutes after EdU injection, animals were sacrificed, and brain samples were harvested and prepared as described in the cell cycle reentry experiment. To measure newborn cell survival, three pulses of EdU (50 μg/g body weight) spaced 24-hours apart were injected intraperitoneally in animals at ∼P90. Animals were sacrificed 30 days later, and brain samples harvested.

##### RNA sample prep and sequencing

All tools, glassware, gloves, and surfaces were cleaned with ethanol and RNaseZAP™(Invitrogen) according to manufacturer guidelines.

Mice were anesthetized using isoflurane. Mice were then placed in a shallow ice trough and rapidly perfused with 10mL ice-cold 1x phosphate buffered saline (PBS) (total time required from induction of anesthesia to end of perfusion was ∼5 min). The brain was immediately removed and transferred to a pre-chilled stainless-steel adult mouse brain matrix placed in ice to make 0.3mm coronal sections (∼2min) using pre-chilled blades. The sections were immediately transferred to ice-cold Hibernate-A in a pre-chilled glass petri dish on ice. Dentate gyrus (DG) was visually identified and dissected out from hippocampal sections to isolate the upper and lower blades along with adjoining regions of the hilus and part of the molecular layer (**See Figure 5A**). The microdissections took no more than 3min per brain. Dissected DG bits were transferred to 1mL RNALater in a microcentrifuge tube placed in ice and were left rotating at 4°C overnight. The following day, tissue was extracted from RNALater solution, flash frozen using dry ice and 100% ethanol and stored at -80°C until RNA extraction.

RNA was extracted from flash frozen DG bits and cultured neurons using the RNeasy Micro Kit (Qiagen) with on-column DNAse treatment following manufacturer guidelines. RNA was poly-A selected and RNA-Seq libraries were prepared using NEBNext Library Prep Kit for Illumina (NEB #E7760). A Miseq shallow sequencing run was performed after library preparation to balance the sample loading process on the deep sequencer. Sequencing was performed using NSQ 500/550 Hi Output KT v2.5 (150 CYS) (Illumina) to obtain 75bp paired end reads. The estimated read depth was ∼33.33 million reads/DG sample.

##### Differential Gene Expression (DEG) Analysis

QC inspection was performed on sequencing data using a combination of FastQC, MultiQC, and Fastx_Screen. Next, adaptor sequences were clipped and low quality and/or nucleotide biased positions in sequences were trimmed using the Trimmomatic; QC inspection of the post clipped and trimmed sequences was repeated. Clipped and trimmed sequences were aligned to the mouse reference genome (mm10) using the HISAT2 tool and QC inspection of the alignments was performed using the RSeqC. Transcript assembly was performed using StringTie which included the discovery of any novel genes:isoforms. QC inspection of the assembly was performed using the gffcompare tool. To enumerate both gene-level and transcript-level expression from the assembly output, IsoformSwitchAnalyzeR package supported in R was used. For differential analysis of the enumerated gene-level expression, commands and functions supported in R were used. Briefly, expression per sample was first within sample normalized via cpm (counts per million) then pedestalled by 2 by taking the log2 transformation. To correct for differences in spread and location, required before testing, cross-sample normalization was performed using Cyclic Lowess. Expression after normalization was inspected by cov-based PCA scatterplot to identify any outliers. Using this 1 sample was identified as an outlier (WT female). This sample was further inspected for coverage using a set of house-keeping genes in mm10 and was found to be 5’ degraded and as such was excluded from further analysis. Other samples were reinspected for coverage and were found to pass QC. Normalization for the gene-level expression was repeated after outlier removal, and sequence batch was globally corrected for via the Limma tool. Using the noise model plot generated from Limma, we identified a mean batch-corrected normalized expression value of 2.25 that was then used to filter genes and transcripts not having at least one sample greater than this value while also flooring values to 2.25 for surviving genes and transcripts if less than this value. These filter-floored values for surviving transcripts were tested for differential expression between WT and V_321_L samples via ANCOVA controlling for sequence batch and biological sex. Dysregulated genes were identified as those having a p-value of <0.05 or <0.1 (colored differently on volcano plot) corrected for batch and/or sex and a minimum fold-change of 1.25 in either direction. Dysregulated genes were then used to perform enrichment analyses using Ingenuity Pathway Analysis (IPA) (Qiagen) and Enrichr (Chen, Tan et al. 2013, Kuleshov, Jones et al. 2016, Xie, Bailey et al. 2021). WGCNA was conducted using the R package (detailed code can be found on the Github link provided below).

##### Network analyses

Schizophrenia-related enriched terms were combined into a single pathway in IPA yielding 67 genes. These 67 genes were selected as seeds and a network was grown using the Build>Grow tool in IPA. The following criteria were set to grow the network: Direct interactions only using the DEG list from the V_321_L vs. WT dataset, using all data sources within the IPA knowledge base, for miRNAs we selected only those that were experimentally observed. For data from cell lines and tissues, we selected only data from CNS cell lines, neuroblastoma cell lines, and neurons using the stringent filter setting to ensure that two nodes will only be connected if the genes are expressed in neural cells. We also excluded “correlation” and “membership” as relationship criteria for connecting nodes leaving behind direct physical or signaling interactions.

Genes from WGCNA modules were used to generate PINs using STRING (Szklarczyk, Kirsch et al. 2023). We adjusted the settings to only include interactions from experiments, databases, and gene fusions. Minimum required interaction score was set to 0.9 (highest stringency) and disconnected nodes were removed.

Networks were exported to Cytoscape and hub gene analysis was performed using the CytoHubba plugin.

##### Statistical analysis

Statistical analysis was performed using Prism (GraphPad). Normality was assessed using Shapiro-Wilkins and Kolmogorov-Smirnov tests. If data failed either test, non-parametric stats were used. p-values were corrected for multiple comparisons as necessary using Bonferroni (parametric) post-hoc test. For Sholl analysis we followed published guidelines to avoid faulty inferences in Sholl analyses (Wilson, Sethi et al. 2017). Sholl analysis data did not pass Shapiro-Wilkins normality test; however, given that measurements were made from multiple neurons from the same animal pooled with neurons across animals, we could not use the Wilcoxon rank sum test given the assumption of independence of variables. To address this, we transformed the data by taking the square-root of the measurements and used a repeated measures two-way ANOVA with a mixed effects model. To control for false discovery rate (FDR) and Type I error inflation, p-values were corrected for multiple comparisons using the original Benjamini-Hochberg FDR procedure.

## DISCLOSURES

The authors have no conflicts of interest to declare.

A version of this manuscript has been posted as a preprint on bioRxiv: https://doi.org/10.1101/2022.08.10.503469

## ACKNOWLEDGEMENTS

This work was supported by the Intramural Research Program of NINDS and by MH087473 (to DAT). We would like to thank Dr. Shaoyu Ge (Stony Brook University, NY) and Dr. Mala Ananth (NINDS) for valuable discussions. We would also like to thank Dr. Abdel Elkahloun and Bayu Sisay (NHGRI sequencing core facility) for providing NGS consultation and services, the NINDS Light Imaging Facility, and Kevin Cravedi (NIMH Rodent Behavior Core facility). Finally, we would like to thank Wendy Akmentin, Taylor Muir and Li Bai for expert technical assistance in data curation. Artwork/Schematics were created with BioRender.com.

## RESOURCE AVAILABILITY

### Lead Contact

Further information and requests for resources and reagents should be directed to and will be fulfilled by Lead Contact, Dr. David Talmage.

### Materials Availability

Plasmids used in this study are available on request.

### Data and Code Availability

The data generated in this publication have been deposited in NCBI’s Gene Expression Omnibus (Edgar, Domrachev and Lash 2002) and are accessible through GEO Series accession number GSE192869 (https://www.ncbi.nlm.nih.gov/geo/query/acc.cgi?acc=GSE192869) and GSE268856 (https://www.ncbi.nlm.nih.gov/geo/query/acc.cgi?acc=GSE268856).

Code is available at https://github.com/RajNINDS/V321L_DG

